# PROST: An Open Toolbox, Harmonised Multi-Lab Dataset, and Meta-Analysis of Ankle-Foot Roll-Over Shape

**DOI:** 10.64898/2026.11.05.764655

**Authors:** Chen Reichsbouscher, Dana Solav

**Affiliations:** Faculty of Mechanical Engineering, Technion Israel Institute of Technology, Haifa, 3200003, Israel

**Author notes:** Corresponding author: Email address (Dana Solav).

**Keywords:** anthropometrics, centre of pressure, effective rocker, gait biomechanics, inter-individual variability, mixed-effects model, prosthetic and orthotic design

## Abstract

The ankle-foot roll-over shape (ROS) is the sagittal-plane path of the centre of pressure (CoP) during the stance phase of walking, expressed in a shank-based coordinate system. It describes the effective rocker geometry that emerges from both the shape and the elastic deformation of the ankle-foot system during walking, and informs prosthetic and orthotic design. Previous studies characterised ROS in small samples with differing laboratory protocols, so they have not systematically assessed inter-individual variability or its relation to anthropometrics.

We developed the open-source Public ROS Toolbox (PROST) for standardised ROS computation, and used it to harmonise marker trajectories, ground reaction forces and CoP data from four public gait datasets collected in independent laboratories. For each leg, we fitted a circular arc to the pooled ROS of all its stance phases. In a one-stage individual-participant data meta-analysis, linear mixed-effects models, with dataset as a fixed effect and participant as a random intercept, related the arc radius and the horizontal coordinate of the circle centre to anthropometrics, and, in the two self-selected-speed datasets, to walking speed.

The harmonised dataset comprises 249 participants and 5,667 stance phases at walking speeds of 0.7–8.5 km/h. Circular arcs fitted the ROS closely (mean RMS radial residual 1.9 mm). The radius had a median of 410.3 mm (interquartile range, IQR, 237.6 mm), and the circle centre lay posterior to the ankle (median −20.6 mm, IQR 49.5 mm). Between-participant differences, quantified by the intraclass correlation coefficient (ICC), accounted for 42% of the variance in log radius (95% CI 25–57%) and 47% of the variance in the horizontal coordinate of the circle centre (95% CI 24–65%). Anthropometrics explained a negligible share of this variance (radius: pseudo-*R*^2^ −0.001, 95% CI −0.03 to 0.12), and height normalisation did not reduce radius variability (robust coefficient of variation 43% with and without normalisation). Faster self-selected walking was associated with a larger radius and a more posterior circle centre.

To our knowledge, this study provides the first harmonised multi-lab ROS dataset. Together with the open-source toolbox, it is publicly available to support reproducible research in gait biomechanics, rehabilitation device design and clinical gait assessment. ROS geometry was participantspecific but was not predicted by age, sex, height or mass. A device ROS scaled from anthropometrics is therefore unlikely to match an individual’s ROS, motivating studies of whether matching it, through device geometry and stiffness, improves gait mechanics.

## 1. Introduction

Human walking gait involves interaction between the foot and the ground across a contact area, where the plantar surface transmits distributed traction, comprising both normal (pressure) and shear components. The centre of pressure (CoP) is defined as the weighted centre of the distributed pressure and represents the point through which the resultant ground reaction force (GRF) vector acts. During stance, the CoP trajectory typically progresses from heel to toe between initial contact (IC) and toe-off (TO). As the body moves forward over the stance leg, the foot-ankle system effectively behaves like a rocker, smoothly governing the centre of mass trajectory with each step [1, 2]. The roll-over shape (ROS) characterises this rocker behaviour and is typically defined from IC to opposite IC (OIC) [1]. Although the CoP continues to advance from OIC to TO, the trailing leg is rapidly unloaded during this terminal double-support phase as weight transfers to the leading leg; thus this phase is typically excluded from ROS characterisation [1]. The ROS provides a compact, functionally meaningful description of human gait mechanics that captures the effective geometry governing forward progression.

The ankle-foot ROS can be quantified by transforming CoP trajectories into a shank-based coordinate system (CS) with origin at the ankle, following the method introduced by Hansen et al. [3, 4], building on Knox [5]. In this approach, CoP measurements obtained from force plates are combined with lower-limb kinematics from motion capture to define the shank-based CS. In this CS, the sagittal-plane CoP trajectory between IC and OIC is often well approximated by a circular arc, which can be fitted using nonlinear least-squares methods [1, 6]. The ROS is thus characterised by circular arc parameters: radius, centre coordinates, and initial and final arc angles.

The rocker-like progression of the foot during stance has long been used to inform the design of assistive devices, even before the ROS concept was formalised. Rocker-bottom therapeutic shoes, commonly prescribed for conditions such as diabetic plantar ulcers, incorporate a rocker sole that can redistribute plantar pressure and alter lower-limb kinematics during gait [7, 8, 9]. Similarly, controlled ankle motion (CAM) boots incorporate a rocker sole, which has been shown to influence plantar pressure distribution and gait mechanics during immobilisation [10, 11]. The ROS provides a framework to quantitatively describe these rocker-based behaviours.

In prosthetic legs, ROS-based analyses have informed the design of ankle-foot systems, including the selection of effective stiffness and geometric parameters to approximate physiological gait [12, 13]. The ROS has also been used as a benchmark for evaluating prosthetic foot performance. Prosthetic designs based on compliant cantilever beams with a constant or monotonically decreasing cross-section have been shown to inadequately reproduce physiological ROS [14]. Moreover, the ROS has been used to evaluate prosthetic alignment with changing heel height [15], and to guide clinical alignment of transtibial prostheses [16]. In many of these applications, a general or population-based ROS was used, with limited consideration of individual variation.

Beyond device design, ROS geometry has also been shown to be functionally significant for walking. Adamczyk et al. attached rigid arc-shaped soles of varying radii and lengths to an ankle immobilisation boot and found that arc geometry affected both the mechanical work performed on the centre of mass and the metabolic cost of walking [17, 18].

Despite this functional significance, the characterisation of ROS parameters remains limited. Prior studies have commonly normalised ROS radius by body height, implicitly assuming proportional scaling. Hansen et al. reported a median ROS height-normalised radius of approximately 0.19–0.22 for able-bodied populations [19, 20]. When computing the knee-ankle-foot ROS, in which the CoP is expressed in a CS aligned with the hip-ankle line rather than the shank, me-dian height-normalised radii of 0.15–0.16 were reported for various walking speeds and carried loads [1, 21]. Although these studies suggested that radius scales approximately with body height, they have not analysed its inter-individual variability.

The horizontal position of the circular arc centre remains poorly characterised and reported, despite its biomechanical importance for both gait interpretation and device design [19, 17]. In observational studies of able-bodied walking, reported values are typically small compared to the radius and tend to be anterior relative to the ankle joint. Median height-normalised values for the knee-ankle-foot ROS range between 0.006–0.011, with inter-individual variability apparent in the reported figures [1, 21]. Although this parameter has been found to vary with walking speed and heel height, these effects were considered of limited clinical significance because the values remain small in relation to body height [4, 1]. Yet, the assumption that small implies negligible does not translate to device design, where the centre horizontal position must be specified. Studies using rigid rocker soles found that even modest fore-aft offsets had pronounced functional consequences: positioning the arc centre directly beneath the tibial axis caused the GRF to pass behind the knee during stance, which required high quadriceps activity to prevent knee buckling, while larger anterior offsets led to discomfort during prolonged walking [17, 18]. A centre horizontal coordinate of 0.076 m [17] and 0.058 m [18] anterior to the tibial axis were ultimately selected through empirical tuning to balance these competing demands. Similarly, Hansen et al. specified the centre horizontal coordinate for the “ideal” ROS as part of a prosthetic alignment hypothesis [16], by drawing on prior literature [5, 22] and clinical trial-and-error, rather than systematic participant-specific analysis [4]. Across these studies, the centre horizontal coordinate was treated as a consequential parameter when poorly set, yet its inter-individual variability and its biomechanical implications have not been systematically characterised.

Despite these important findings and advances, several limitations remain, which share a common thread: ROS parameters have been studied in relatively small, heterogeneous cohorts, and inter-individual variability has not been systematically linked to measurable anthropometric factors. The common practice of normalising ROS parameters by body height rests on an implicit assumption of proportional scaling that has not been rigorously validated. Critically, the determinants of the centre horizontal coordinate remain insufficiently understood.

To address these gaps, we compiled multiple publicly available datasets containing motion capture and force-plate measurements into a unified database of *N̄ _p_* = 254 participants. We computed shank-based CoP trajectories and estimated ROS circular arc parameters for each participant. We then characterised population-level distributions and examined relationships with anthropometric variables. For the radius and the centre horizontal coordinate, we additionally quantified inter-participant variability, and its association with anthropometrics and walking speed, using linear mixed-effects models. This large-sample analysis provides a more robust assessment of inter-individual variability, evaluates the validity of ROS anthropometric scaling, and provides population-level reference distributions for both radius and centre horizontal position. The unified database and computational toolbox accompanying this work facilitate straightforward replication and extension of these analyses. Taken together, these contributions advance the biomechanical understanding of human gait and provide empirical evidence to inform the design and personalisation of orthotic and prosthetic devices.

## 2. Methods

The *Public ROS Toolbox* (*PROST*, v1.0.0), accessible at https://github.com/SolavLab/ PROST, was developed as an end-to-end framework for reproducible individual- and populationscale analysis of ankle-foot ROS parameters. Implemented in MATLAB R2024b (The Math-Works, Inc., Natick, MA, USA) with the Optimization, Statistics and Machine Learning, and Signal Processing Toolboxes, it is an open-source toolbox for pooling gait biomechanics data from multiple publicly available datasets, computing ROSs, fitting circular arc parameters, and analysing their relationships with anthropometric variables. The overall workflow consists of preprocessing raw individual-level walking trials from four open datasets, harmonising datasetspecific variables and marker definitions, standardising measurements into a common spatial and temporal representation, and pooling the resulting data into a unified database. We then computed the right and left ROS for each participant and the corresponding circular arc parameters, which were subsequently used in an individual participant data (IPD) meta-analysis.

### 2.1. Dataset selection

To construct the database, we defined four inclusion criteria: datasets must (1) contain 3D marker trajectory data of the lower limbs (including knee and ankle) and force-plate measurements of GRF and CoP for healthy participants during walking; (2) include participant-level anthropometric data; (3) contain at least one trial with a complete stance phase; and (4) provide or allow the derivation of the event times of IC, OIC and TO for each step.

Using these criteria, we conducted a systematic search of publicly available motion capture repositories and identified four datasets [23, 24, 25, 26], comprising a total of *N̄ _p_* = 254 healthy participants. The main characteristics of the datasets are summarised in Table 1.

**Table 1:** Summary of datasets included in the database. M/F indicates male and female participants. Numeric values are reported as mean ± standard deviation. Dataset 1’s speed categories are nominal (metronome cued), while dataset 2’s speed protocols were only confirmed for the first trial of each condition.

|  | Dataset 1 | Dataset 2 | Dataset 3 | Dataset 4 |
|---|---|---|---|---|
| Manuscript | Moreira et al., 2021 [23] | Schreiber et al., 2019 [24] | Lencioni et al., 2019 [25] | Van Criekinge et al., 2023 [26] |
| Participants | 16 | 50 | 50 | 138 |
| Sex | 8 M, 8 F | 26 M, 24 F | 25 M, 25 F | 65 M, 73 F |
| Age [years] | 23.8 ± 2.1 | 37.0 ± 13.6 | 27.0 ± 17.6 | 51.2 ± 19.6 |
| Height [m] | 1.69 ± 0.11 | 1.74 ± 0.09 | 1.60 ± 0.20 | 1.68 ± 0.10 |
| Body mass [kg] | 67.5 ± 11.1 | 71.2 ± 12.3 | 56.7 ± 19.8 | 74.4 ± 15.1 |
| Footwear condition | Shod | Barefoot | Barefoot | Barefoot |
| No. force plates | 6 | 2 | 2 | 4 |
| Speed categories [km/hr] (reported on) | • 1.0<br>• 1.5<br>• 2.0<br>• 2.5<br>• 3.0<br>• 3.5<br>• 4.0 | • 0 - 1.44<br>• 1.44 - 2.88<br>• 2.88 - 4.32<br>• Self-selected (spontaneous)<br>• Self-selected (fast) | • 5 self-selected.<br>• 5 progressively increasing trials.<br>• 5 progressively decreasing trials. | Self-selected |

### 2.2. Data Preprocessing

All datasets were preprocessed to ensure consistency and reliability of the extracted gait parameters. Walking trials were stored as complete trial recordings for reference and traceability. Step-level information, including IC, OIC and TO event times, quality classification (good or bad), step number and force plate number, was stored separately for each step. Good-quality steps were defined as those in which a single foot maintained full contact with a single force plate throughout the stance phase. This quality classification, along with the stored event times, was subsequently used during harmonisation to segment trials into individual steps, retaining only good-quality steps for analysis.

Anthropometric data were stored for each participant as described in Table 2, along with the participant’s footwear condition (shod or barefoot; see Table 1).

**Table 2:** Anthropometric parameters, their units, and their source or calculation method.

| Parameter | Unit/Category | Source or calculation Method |
|---|---|---|
| Age | years | Reported |
| Sex | Male/Female | Reported |
| Height | m | Reported |
| Mass | kg | Reported |
| Leg length | m | Distance between the anterior-superior iliac spine and lateral ankle markers. |
| Thigh length | m | Distance between the anterior-superior iliac spine and lateral knee markers. |
| Shank length | m | Distance between the lateral knee and lateral ankle markers. |

During preprocessing, several issues were identified that precluded direct harmonisation and required additional processing steps. Dataset 3 [25] lacked OIC event times, and datasets 2 and 3 [25, 24] were missing knee markers. These issues were addressed by estimating OIC events from heel marker heights and reconstructing virtual knee markers using available femoral and tibial markers, as detailed in Appendix A and Appendix B, respectively.

### 2.3. Data Harmonisation and Standardisation

After preprocessing, each dataset was reorganised into a standardised hierarchical structure and stored as individual .mat files. To facilitate harmonisation across datasets, measurements were standardised using common CS, temporal, and naming conventions. A schematic of the main fields is shown in Figure 1. The standard structure comprises the following top-level fields, with full formatting specifications provided in the toolbox documentation:

- Article data: metadata about the source article, including the title, authors, year of publication, and URLs of the article and data files.
- Markers guide: a reference mapping the harmonised marker names to the original marker names or measurement locations. This ensures consistent naming across datasets to facilitate automated processing. When knee markers were estimated from other markers, the guide lists the source markers and labels the knee marker as “virtual”.
- Lab system: a description of the laboratory setup, including the CS used in the dataset and the number of force plates in the experiment.
- Experiment protocol: a description of the experimental protocol, including the walking speed categories and the number of participants.
- Data: the field containing participant-level characteristics and measurement data.

**Figure 1:**
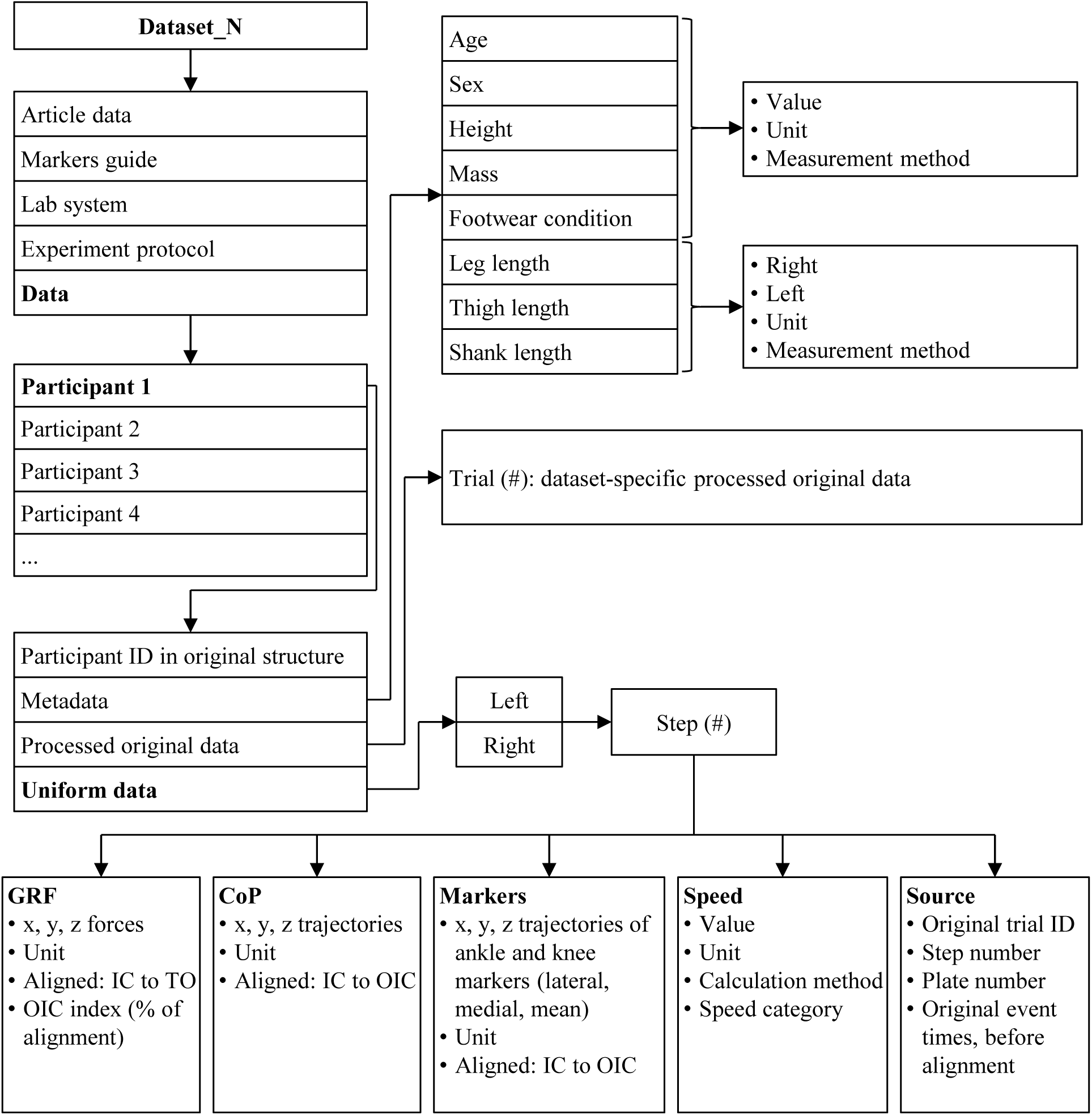
Schematic of the harmonised dataset structure. Steps are organised separately for the left and right legs, which may contain different numbers of steps. “Dataset N” refers to the object name of the specific dataset where *N* = 1, 2, 3, 4 in our case. GRF: ground reaction force; CoP: centre of pressure; IC: initial contact; OIC: opposite initial contact; TO: toe-off; ID: identifier.

The Data field is organised hierarchically by participant and includes anthropometric data, footwear condition, the pre-standardisation trial data (retained from preprocessing; see section 2.2), and the standardised measurement data used for ROS computation and statistical analyses.

The anthropometric data for each participant include parameters that were either reported or estimated from recorded marker positions. The procedure for obtaining each parameter is described in Table 2, along with the corresponding units. Leg, thigh, and shank lengths were estimated for each leg from specific marker distances measured during the standing trial, or, when a standing trial was unavailable, estimated as the median across trials of the per-trial median marker distances. These estimates were used for all datasets, including those in which the values were originally provided, to ensure consistency across datasets regardless of original reporting format.

The standardised data are divided into left and right steps, with each step stored individually, even if the original trials include multiple steps. Only the steps that passed the quality screening are retained. Marker trajectories consist of both lateral and medial markers (when available), as well as their mean trajectory. When only lateral markers are present, only lateral trajectories are stored.

Within each step, measurement data were further standardised into a common spatial and temporal representation. All data were first transformed into a standardised laboratory CS defined as follows: **x̂** represents the forward walking direction, **ẑ** is the vertical axis (pointing up), and **ŷ** completes a right-handed CS (pointing to the left). The CoP and marker trajectories were then temporally aligned to 101 points (0–100%) between IC and OIC event times. This standardisation facilitates automated transformation of the CoP into the shank-based CS using the knee-ankle marker trajectories, allowing for automation of the ROS computation.

Additional stored parameters include GRF measurements temporally aligned to stance-phase percentage, the average walking speed, and step-level information from the original dataset, including the original trial ID to which each step belongs and the event times and frame indices prior to temporal alignment. Table 3 outlines the structure of the step data.

**Table 3:** Standardised data parameters stored for each participant, with corresponding time intervals. All trajectories and forces were stored in the laboratory coordinate system (CS).

| Parameter | Unit | Time interval |
|---|---|---|
| CoP | mm | IC to OIC |
| GRF | N | IC to TO |
| Ankle Markers | mm | IC to OIC |
| Knee Markers | mm | IC to OIC |
| Speed | km/hr | IC to TO |

### 2.4. ROS Analysis Framework

After harmonising and standardising the data, we developed a unified MATLAB class named *PROSTAnalyzer*, which serves as the central interface for data access, integration, and analysis. The class imports each standardised dataset and stores it in a data records structure that preserves the hierarchical organisation of datasets and participants.

*PROSTAnalyzer* maintains two primary analysis tables: (i) a dataset-level summary table that describes cohort characteristics (see Table 1), and (ii) a leg-level IPD analysis table that consolidates anthropometrics, footwear condition, and ROS-derived circular arc parameters, computed as described in section 2.4.1. This table forms the basis for all subsequent statistical analyses and visualisations.

#### 2.4.1. ROS Computation

To compute each leg’s ROS, we transformed the temporally aligned CoP trajectories of each step into a shank-based CS. This CS, defined using the lateral ankle and knee marker trajectories (medial markers were not available in all datasets), was constructed following the formulation of Hansen et al. [20]. The CS consists of three orthonormal vectors **û**, **v̂**, **ŵ** , with origin at the ankle. Let **p**_ankle_ and **p**_knee_ denote the ankle and knee coordinates in the laboratory CS (**x̂**, **ŷ**, **ẑ**). We define,

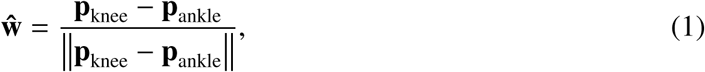

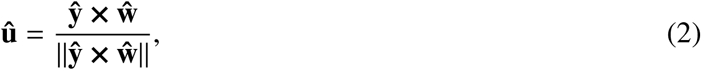

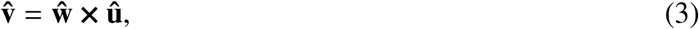

which yields an orthonormal, right-handed CS aligned with the shank, as illustrated in Figure 2. An example of CoP and marker trajectories in the laboratory CS and after transformation to the shank-based CS are shown in Figure 3.

**Figure 2:**
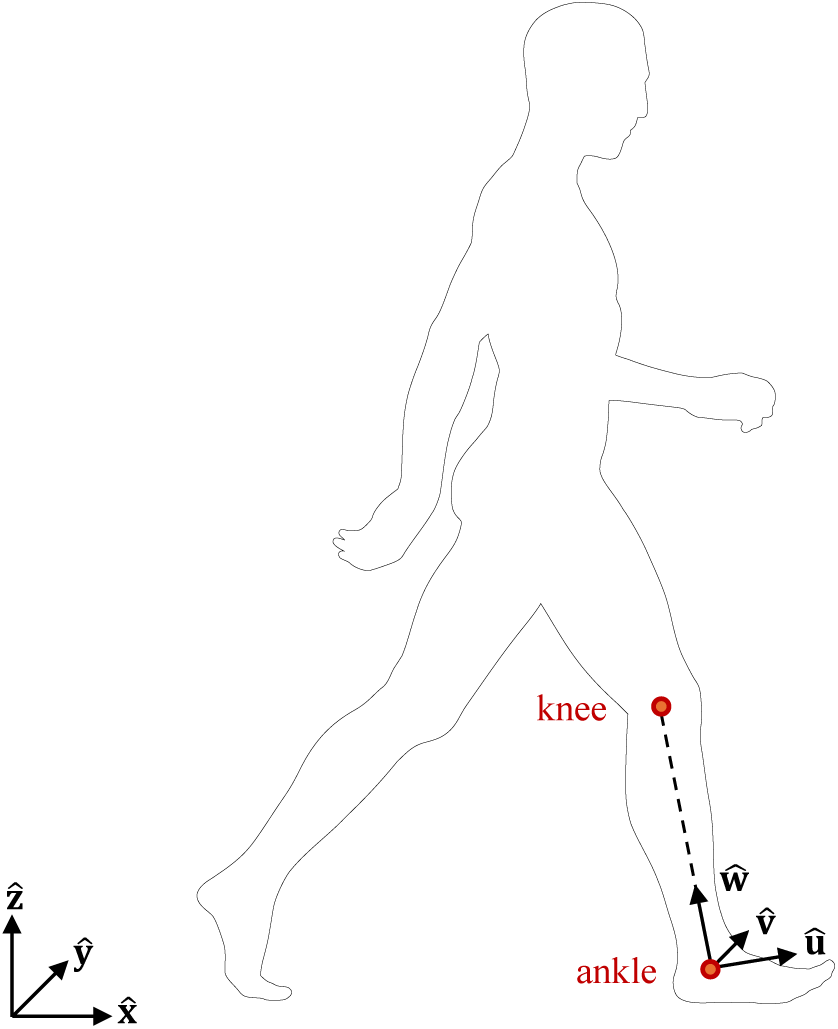
Illustration of the shank-based coordinate system (CS) (**û**, **v̂**, **ŵ** ). The origin is set at the ankle. **ŵ** is a unit vector pointing from the ankle to the knee. **û** is the normalised cross product **û** = (**ŷ** × **ŵ**)||**ŷ** × **ŵ** ||. **v̂** completes a right-handed CS: **v̂** = **ŵ** × **û**. **x̂**, **ŷ**, **ẑ** are the axes of the laboratory CS. Figure overlay adapted from “Silhouette of a walking man” by T.M.M. Aumala, Wikimedia Commons, 2021 (CC0).

**Figure 3:**
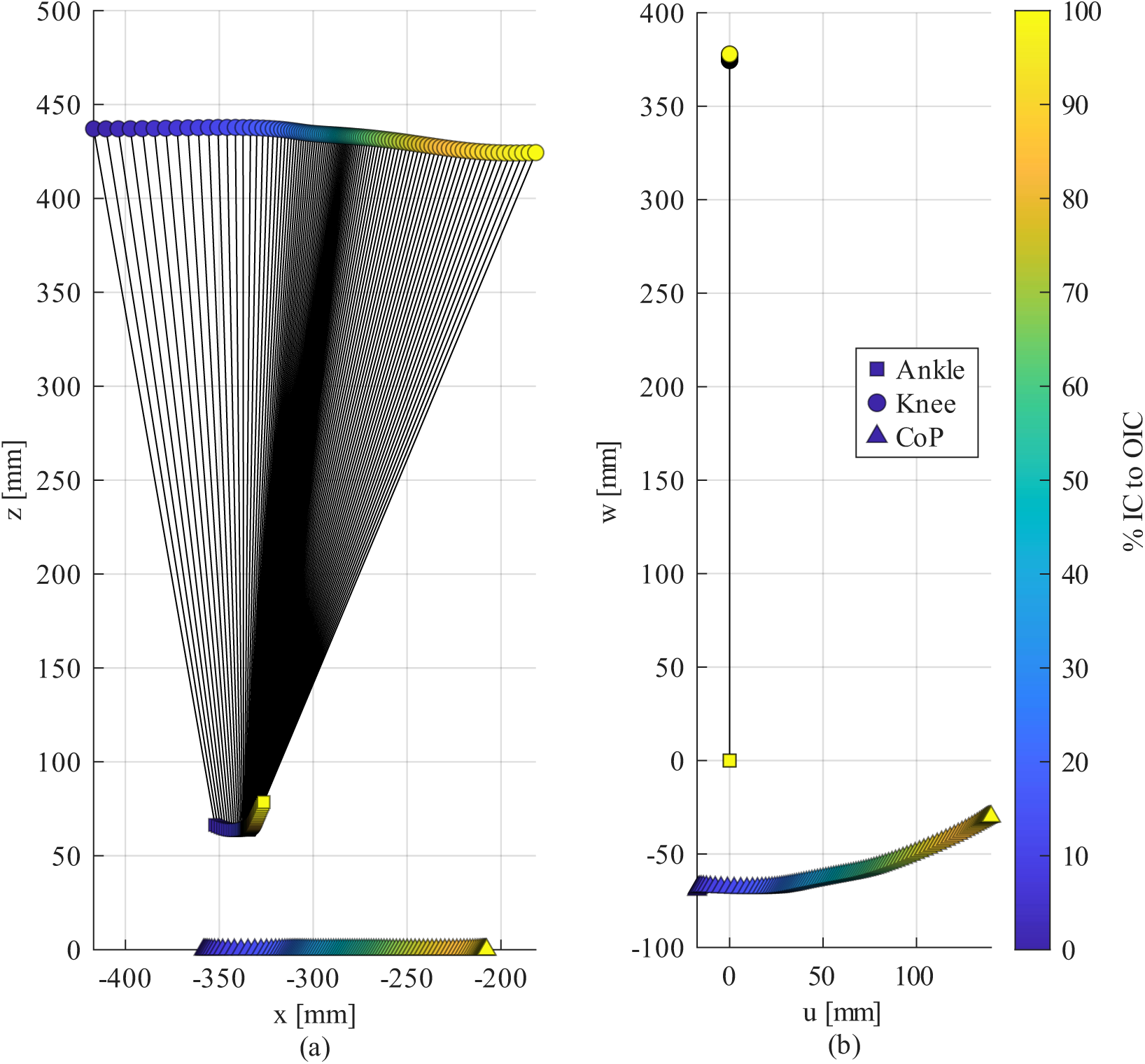
Roll-over shape calculation by coordinate system transformation of marker trajectories and centre of pressure. For one left-leg step of participant 7 in Dataset 2 [24], ankle (squares), knee (circles), and centre of pressure (CoP; triangles) are shown (a) in the laboratory coordinate system (CS) (**x̂**–**ẑ** plane) and (b) after transformation to the shank-based CS (**û**–**ŵ** plane). Black lines connecting the ankle and knee markers illustrate the shank orientation. Colour denotes normalised time from initial contact (IC, 0%) to opposite IC (OIC, 100%).

ROS fitting was performed at the leg level using all retained steps. For each leg, we pooled all shank-based CoP coordinates into a single point cloud in the **û** − **ŵ** plane. We then fitted a circular arc to this point cloud, computing the best-fit radius *r*, horizontal and vertical coordinates of the circle centre (*u_c_, w_c_*), and initial and final angles of the circular arc relative to the −**ŵ** direction. The arc fitting followed the method in Bapat et al. [6]. We first solved the reduced-order linear least-squares problem [27]. Then, using an initial guess derived from this solution, we computed a nonlinear total least-squares fit of the circular arc with MATLAB’s fminunc optimisation function, which uses a quasi-Newton algorithm. The objective function was the sum of squared radial residuals between the CoP points and the fitted circle:

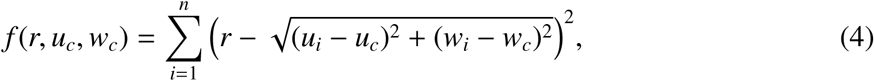

where *n* is the number of data points, and (*u_i_, w_i_*) are the coordinates of the CoP data points in the shank-based CS. We then calculated the initial and final circular arc angles from the circle parameters and the minimum and maximum values of *u_i_*.

The root-mean-square (RMS) radial residual of each leg, evaluated at the best-fit parameters *r*^∗^*, u_c_*^∗^*, w_c_*^∗^, was stored in the IPD analysis table as a measure of goodness of fit, as:

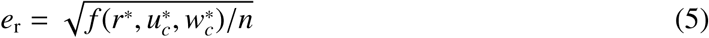

### 2.5. Statistical Analysis

All statistical analyses were performed at the leg level in MATLAB R2024b (Statistics and Machine Learning Toolbox) and were restricted to two circular arc parameters: the radius *r*, the parameter frequently reported in previous studies, and the centre horizontal coordinate *u_c_*, which determines the fore-aft position of the arc relative to the ankle. Height-normalised parameters were obtained by dividing the radius and the centre horizontal coordinate by participant height, and the height-normalised radius is denoted *R_H_*. Both parameters were modelled without normalisation, the radius in millimetres and on the log scale and the centre horizontal coordinate in millimetres, with height entering the covariate models as a covariate (sections 2.5.2 and 2.5.3). All hypothesis tests were two-sided with a significance level of *α* = 0.05. The pairwise anthropometric association analyses (Appendix C) were exploratory, and their p-values are reported without correction for multiple comparisons; in the covariate models, the anthropometric effects were estimated jointly and adjusted for dataset (section 2.5.3).

#### 2.5.1. Distributions and Pairwise Associations

The distributions of *e*_r_ and the circular arc parameters were summarised by their mean and standard deviation (SD), median and interquartile range (IQR), skewness, and kurtosis (non-excess, where a normal distribution has a kurtosis of 3). Normality of *e*_r_ was assessed with the Anderson-Darling test, both pooled and per dataset. For the radius, relative variability was quantified using the quartile-based robust coefficient of variation (RCV*_Q_* = 0.75 × IQR*/*median) [28], with the conventional coefficient of variation (CV = SD/mean) reported for comparison. A t location-scale distribution was fitted by maximum likelihood to the radius and the centre horizontal coordinate, raw and height-normalised, for visual comparison with their histograms.

Pooled pairwise associations between the raw and height-normalised circular arc parameters and the anthropometric variables (Spearman’s *ρ* and *R*^2^ for numeric variables; Wilcoxon ranksum tests with the rank-biserial correlation *r_rb_*for sex and footwear condition) were computed descriptively, treating each leg as an independent observation; their methods and results are given in Appendix C.

#### 2.5.2. Linear mixed-effects model

Because the data are nested (datasets → participants → legs), observations are not independent and identically distributed, and the pooled median and IQR cannot isolate inter-participant variability [29, 30]. We therefore modelled *r* with a linear mixed-effects model, with dataset as a fixed effect and participant (nested within dataset) as a random intercept:

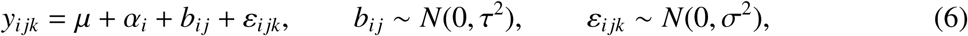

where *y_i jk_* is the response of leg *k* of participant *j* in dataset *i*, *µ* is the grand mean, *α_i_*is the effect of dataset *i* under effects (sum-to-zero) coding 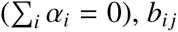 is the participant random intercept, and *ε_i jk_*is the residual. The model was fitted by restricted maximum likelihood (REML). Legs with *R_H_ >* 1, i.e. a fitted radius longer than body height, were excluded before fitting. For a nearly flat CoP path the arc span *θ* is small and the fitted radius is ill-conditioned, since a small change in path curvature produces a large change in *r*. For an arc with chord length *L*, *r* = *L/*(2 sin(*θ/*2)) exactly; because the height-normalised chord *L_H_* varied little across legs (median 0.091, IQR 0.016) compared with *R_H_*, the radius was governed by the arc span, and *R_H_* = 1 corresponds to *θ* ≈ 5^◦^ for a typical chord. The *R_H_* criterion and the equivalent arc-span criterion identified the same 14 legs. The derivation and a sensitivity analysis of the threshold are given in Appendix E. The model was first fitted to *r* and, because the model diagnostics indicated non-normality and heteroscedasticity, refitted to log *r*.

The same model was fitted to *u_c_*, on the same legs. Because *u_c_* is signed and takes values on both sides of the ankle, it was modelled in millimetres without transformation, and height entered its covariate models as a covariate rather than as a divisor (section 2.5.3). The *R_H_ >* 1 exclusion applies to *u_c_* as well: for a nearly flat arc the centre lies far from the CoP path, and its horizontal position is as poorly determined as the radius.

The proportion of total variance attributable to inter-participant differences was quantified by the intraclass correlation coefficient (ICC), also known as repeatability [30, 31]:

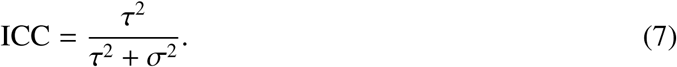

For *u_c_*, which is modelled on its original scale, Equation 7 applies directly. For the model fitted to log *r*, the log-scale ICC, denoted ICC*_log_*, was back-transformed to the original *r* scale assuming log-normality, using the law of total variance:

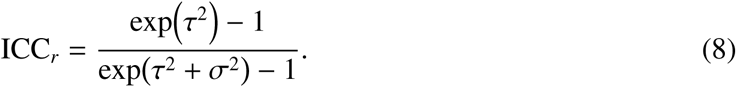

#### 2.5.3. Covariate models

To estimate how much of the between-participant variance was associated with anthropometrics and walking speed, covariates were added as fixed effects to the model of Equation 6:

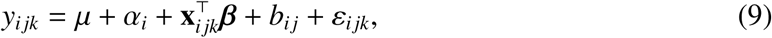

where **x***_i jk_* is one covariate set and *β* its coefficients. Two sets were fitted, each separately: the anthropometric set (age, sex, log *H* and log *M*, with *M* the body mass), and walking speed (*v*^∗^, defined below). Continuous covariates were centred within dataset. Height and mass entered on the log scale, so their coefficients are elasticities of *r*, i.e. the relative change in *r* per relative change in *H* or *M*, as expected for allometric scaling with body size; for height, 1 corresponds to proportional scaling. In a sensitivity analysis, log leg length replaced log *H*. Leg side was fitted as a separate covariate set, as a modelling check: because both legs of a participant share *b_i j_*, side varies within participants and can only capture a right−left shift common to all participants, so it tests whether the two legs can be pooled rather than explaining between-participant variance. Sex and side were entered with effects coding, and their effects are reported as the difference between the two categories. Walking speed was entered as each leg’s mean measured step speed in dimensionless form, 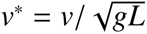, with *L* the leg length. Speed was analysed only in Datasets 3 and 4. In Datasets 1 and 2 each leg pooled several cued speed conditions, in mixes that differed between legs, so a leg-mean speed reflects the protocol and condition mix rather than a participant’s walking speed (Table 1; Appendix D).

The anthropometric and speed sets were assessed by the ICC of the covariate model and by the level-2 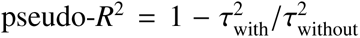 [32], the share of the between-participant variance the set accounts for, where 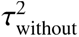 is that of the base model fitted on the same legs. Because the anthropometric covariates are participant-level, and *v*^∗^ almost entirely so, they can only explain between-participant variance, so pseudo-*R*^2^ measures the share of *τ*^2^ they account for; small negative values reflect sampling variability around zero. Whether the covariate effects differed between datasets was assessed by adding covariate-by-dataset interactions. Proportional scaling of the radius with height was assessed by regressing log *r* on log *H* in the same model structure; a slope of 1 corresponds to *r* ∝ *H*, which is what height normalisation assumes. The analyses were repeated in Dataset 4 alone. The covariate models of *u_c_* used the same covariate sets, baselines and speed restriction, with height and mass entered linearly (centred *H* and *M*) instead of on the log scale, since a signed response is not log-transformed and an elasticity is not defined for it. No proportional-scaling test was performed for *u_c_*: proportionality (*u_c_* = *kH*) constrains the intercept rather than the slope, and the intercept is absorbed by the dataset effects. CIs of the coefficients and of pseudo-*R*^2^ were obtained from the case bootstrap described below, refitting both the covariate model and its baseline in each resample. Maximum-likelihood LRTs between nested models are reported as supporting evidence only, since their *χ*^2^ reference assumes the normal, homoscedastic errors that were not met.

#### 2.5.4. Model diagnostics and inference

Model assumptions were assessed using Anderson-Darling normality tests on the pooled response, the Pearson residuals and the predicted random effects, and a Brown-Forsythe test of equal residual variance across datasets, supplemented by visual inspection of residual-versus-fitted and QQ plots. The participant random intercept was tested using a likelihood ratio test (LRT) comparing REML fits with and without the random intercept and with identical fixed effects [33]. Since the null value *τ*^2^ = 0 lies on the boundary of the parameter space, the *χ*^2^ reference distribution is conservative for this test [33]. As a sensitivity analysis, the ICC of each model was recomputed with each participant left out in turn. For the covariate models, the same checks were supplemented by plots of the marginal residuals against each continuous covariate with a quadratic term added to test linearity, variance inflation factors, and pseudo-*R*^2^ recomputed with each participant left out in turn.

Because the normality assumptions were not met, standard errors (SE) and confidence intervals (CI) were obtained by a stratified cluster (case) bootstrap [34] rather than by model-based Wald approximations. The bootstrapping seed used was 20261006. In each of 5000 resamples, participants were drawn with replacement within each dataset, retaining all legs of each sampled participant, and the model was refitted. The SE was taken as the SD of the bootstrap distribution, and the 95% CI as its 2.5th–97.5th percentile interval. Because the pooled model assumes a residual variance common to all datasets, the ICC was also estimated within each dataset, by fitting the same response with an intercept and the participant random intercept only, to each dataset separately. Their CIs were obtained from the same stratified bootstrap, refitting the four dataset-specific models in each resample, so that the datasets were resampled independently of one another.

To address residual heteroscedasticity, observation-level weights were considered. The linear mixed-effects framework allows a heterogeneous residual variance with inverse-variance weights [33, 35]. Two candidate precision measures were evaluated: *e*_r_ and the number of steps used to fit the arc (*n*_steps_). Because the two may not be independent, the association between them was first as-sessed using Spearman’s *ρ* and a log-log power-law fit. Because *n*_steps_ also differs strongly between datasets (see Table 4), we tested whether its relation with precision was confounded by dataset ID. Legs were binned by *n*_steps_ and a power law was fitted between the within-bin variance of log *r* and the bin-midpoint *n*_steps_ on a log-log scale, before and after subtracting dataset means (bins with fewer than 15 legs were excluded from the dataset-adjusted fit). The dataset-adjusted relation was additionally fitted by linear regression weighted by bin size.

**Table 4:** Dataset composition. Counts of participants, legs, and steps are reported as valid / total. Steps per participant are reported as mean ± SD (standard deviation). Walking speed is the measured speed of the valid steps, reported as mean ± SD (range); its distribution per dataset and speed condition is shown in Figure D.15.

|  | Dataset 1 [23] | Dataset 2 [24] | Dataset 3 [25] | Dataset 4 [26] | **Total** |
|---|---|---|---|---|---|
| Participants | 14/16 | 49/50 | 50/50 | 136/138 | **249**/**254** |
| Legs | 28/32 | 98/100 | 76/78 | 265/268 | **467**/**478** |
| Steps | 2683/3153 | 1522/2286 | 812/844 | 650/1350 | **5667**/**7633** |
| Steps per Participant | 191.6 ± 28.2 | 31.1 ± 5.8 | 16.2 ± 9.1 | 4.8 ± 2.0 | − |
| Walking speed [km/h] | 2.56 ± 1.03 (0.86–4.87) | 3.60 ± 1.39 (0.66–8.45) | 4.41 ± 1.31 (1.05–7.80) | 4.28 ± 0.59 (2.61–6.61) | − |

## 3. Results

The following results characterise the ROS radius and centre horizontal coordinate, computed from the lateral markers of the harmonised dataset (see sections 2.4.1 and 2.5).

### 3.1. Database Overview and Participant Flow

After data harmonisation and step-level screening, the final dataset comprised *N_s_* = 5, 667 valid stance phases from *N_L_* = 467 trailing legs of *N_p_* = 249 out of *N̄ _p_* = 254 participants. Three participants were excluded due to missing marker data and another two participants due to inconsistent marker trajectories that produced non-physiological shank-based CoP paths. In addition, step-level screening after harmonisation removed trials with kinematic inconsistencies, supplementing the preprocessing quality screening described in section 2.2. Table 4 details the valid participants and steps used in each dataset.

Dataset contributions were uneven across sources, with Dataset 1 [23] providing the highest number of steps per participant (191.6 ± 28.2), compared to substantially lower values in other datasets. Additionally, Dataset 4 contributed the largest number of valid participants (136), considerably more than the other datasets (see Table 4).

### 3.2. Model Fit and Validation

The circular arc model provided an accurate fit to the ROS across participants. The mean ± SD of *e*_r_ was 1.9 ± 1.0 mm, corresponding to less than 0.5% of both the mean (524.3 mm) and median (410.3 mm) fitted radii. Figure 4a shows a histogram of *e*_r_ across all datasets.

**Figure 4:**
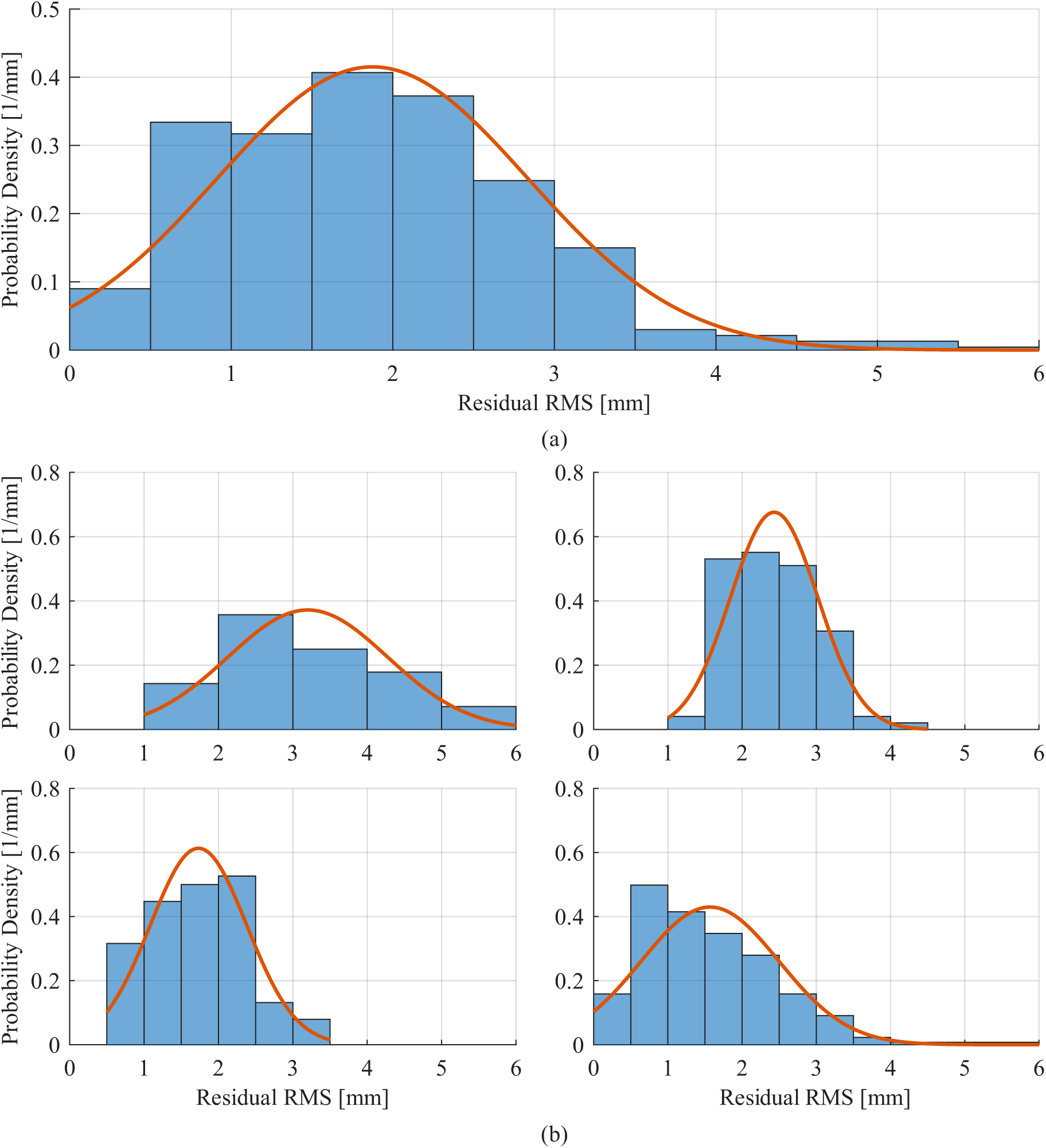
Distribution of the root-mean-square (RMS) radial residual *e*_r_ across all participant-leg circular arc fits: (a) the entire database (*N_L_* = 467 legs), and (b) each dataset separately – Dataset 1 [23] (top left, *N_L_* = 28), Dataset 2 [24] (top right, *N_L_* = 98), Dataset 3 [25] (bottom left, *N_L_* = 76) and Dataset 4 [26] (bottom right, *N_L_* = 265). The fitted normal distribution (red) is compared with the histogram of *e*_r_, normalised as a probability density (blue). The mean ± standard deviation of *e*_r_ is 1.9 ± 1.0 mm and the median (interquartile range) is 1.8 (1.3) mm.

Analysis of the pooled *e*_r_ indicated a deviation from normality, where an Anderson-Darling (AD) test rejected normality (*A*^2^ = 2.03, *p <* 0.001). When stratified by dataset (see Figure 4b), Dataset 4 [26] exhibited deviation from normality (*A*^2^ = 3.28, *p <* 0.001), whereas datasets 1, 2, and 3 (*A*^2^ = 0.55, *p* = 0.14, *A*^2^ = 0.64, *p* = 0.091, *A*^2^ = 0.48, *p* = 0.23, respectively) did not, although the small sample of Dataset 1 (*N* = 28 legs) gives the test limited power.

Representative examples of fitted ROS curves are shown in Figure 5. They demonstrate close agreement between measured CoP trajectories and the fitted circular arc, with *e*_r_ equal to 1.66 mm and 1.29 mm for the right and left legs, respectively.

**Figure 5:**
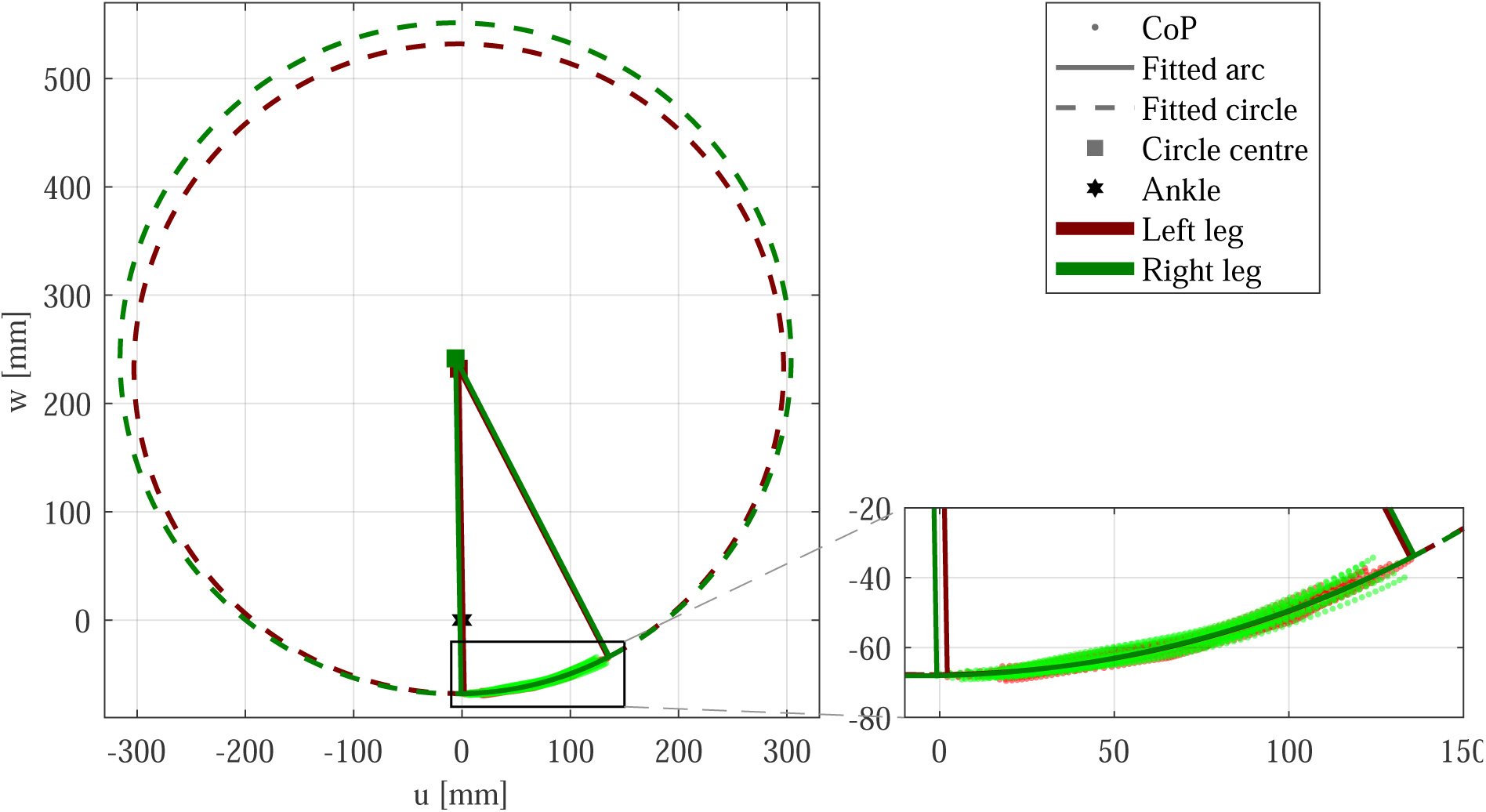
Circular arc fits to the roll-over shape of both legs of a representative participant (participant 7, Dataset 2 [24]), using all valid steps. Centre of pressure (CoP) points from every step are shown in the shank-based coordinate system (*u*–*w* plane) as red (left leg) and green (right leg) dots, together with the fitted circular arc (solid line), the full fitted circle (dashed line) and the circle centre (square); the two lines from each centre mark the start and end of the arc, and the star at the origin marks the ankle. The rectangle outlines the region enlarged on the right. The left-leg arc has a radius of *r* = 299.9 mm, a centre coordinate at (*u_c_, w_c_*) = (−3.0, 232.1) mm and a root-mean-square radial residual of *e*_r_ = 1.29 mm; the right-leg arc has a radius of *r* = 309.8 mm, a centre coordinate at (*u_c_, w_c_*) = (−5.9, 241.7) mm and *e*_r_ = 1.66 mm.

### 3.3. Distribution of ROS Parameters

ROS parameters exhibited non-Gaussian, heavy-tailed distributions as shown in Figure 6a– b, with high skewness and kurtosis (Table 5). The heavy tails were present within Datasets 2– 4 and not only in the pooled sample, whereas Dataset 1 was close to Gaussian (kurtosis near 3). The negative pooled skewness of the centre horizontal coordinate indicates an asymmetry in which more values are located on the posterior side of the ankle reference frame; Dataset 3 was the exception, with positive skewness. Variability is therefore summarised using median (IQR), pooled and per dataset, in Table 5. Mean ± SD is presented for comparison as: 524.3 ± 410.9 mm for the radius and -39.8 ± 103.6 mm for the centre horizontal coordinate.

**Figure 6:**
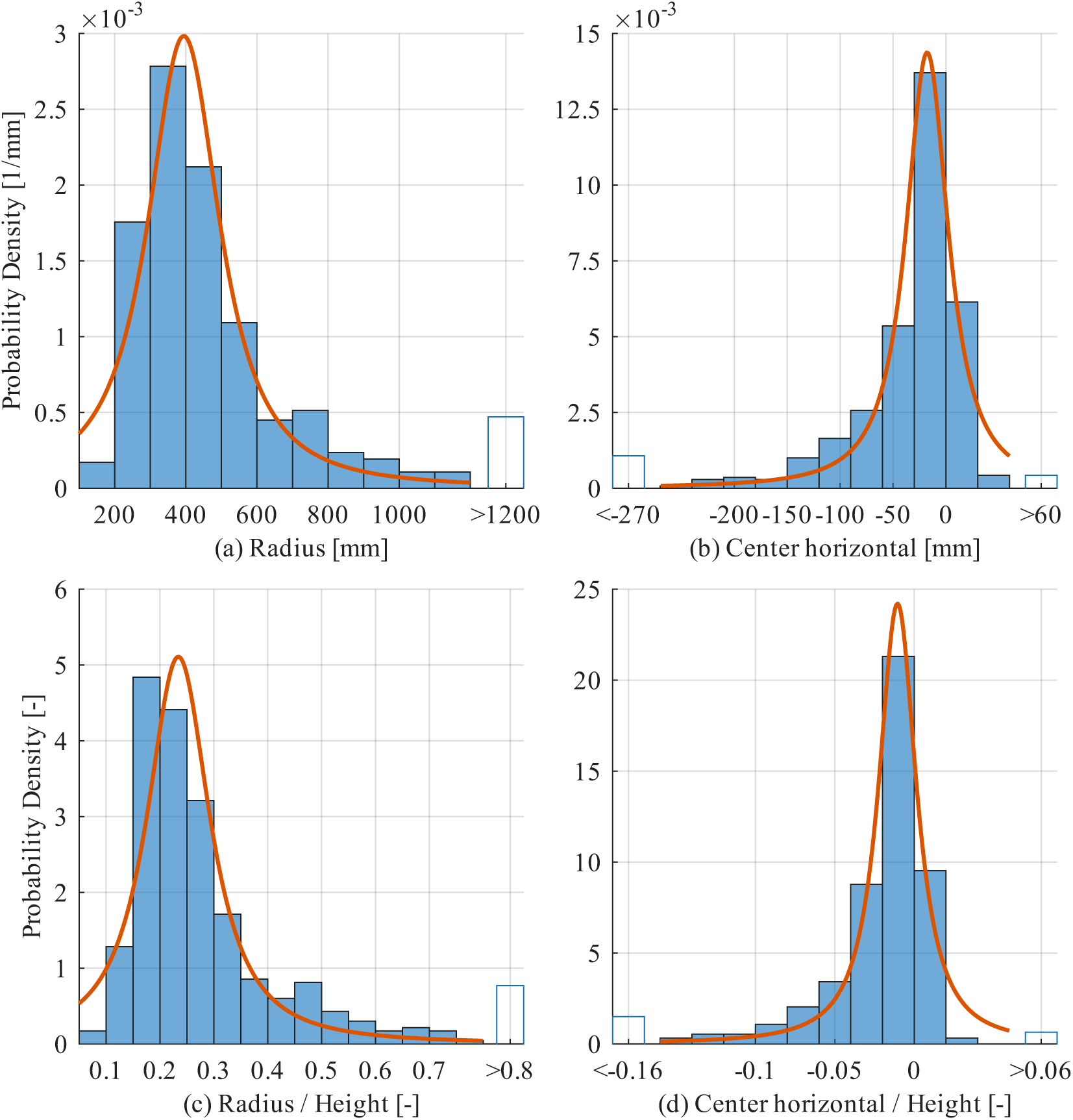
Distributions of the fitted circular-arc parameters over all participant legs (*N_L_* = 467): (a) the radius *r* and (b) the centre horizontal coordinate *u_c_* (mm, positive anterior to the ankle), and (c) *r/H* and (d) *u_c_/H*, normalised by participant height *H*. Histograms (blue) are normalised to unit area and shown with the maximum-likelihood t location-scale distribution (red). Each horizontal axis spans the shortest interval containing at least 95% of the legs; the remaining legs fall in hollow overflow bins, labelled by their range, whose area equals their fraction of legs ((a) 22 legs above; (b) 15 below, 6 above; (c) 18 above; (d) 14 below, 6 above).

**Table 5:**
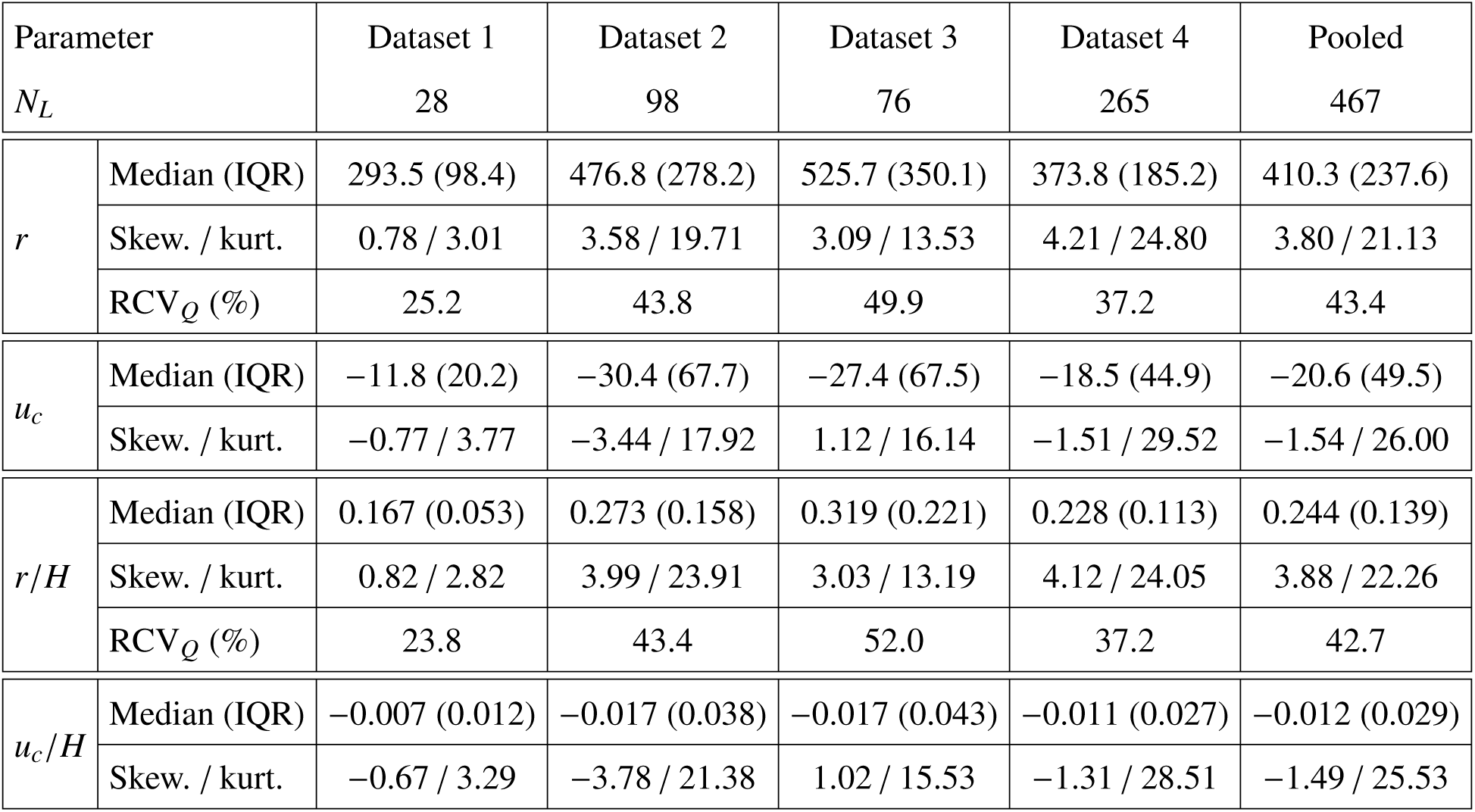
Median (IQR - interquartile range), skewness and kurtosis of the circular arc parameters per dataset and pooled. The quartile-based robust coefficient of variation, RCV*_Q_* = 0.75 IQR*/*median (%), is also given for the radius. Kurtosis is non-excess (3 for a normal distribution); skewness and kurtosis are descriptive and unstable for small samples. *r*: radius (mm); *u_c_*: centre horizontal coordinate (mm), positive anterior to the ankle; *H*: participant height; *N_L_*: number of legs.

| Parameter |  | Dataset 1 | Dataset 2 | Dataset 3 | Dataset 4 | Pooled |
| *N*<sub>*L*</sub> |  | 28 | 98 | 76 | 265 | 467 |
|---|---|---|---|---|---|---|
| *r* | Median (IQR) | 293.5 (98.4) | 476.8 (278.2) | 525.7 (350.1) | 373.8 (185.2) | 410.3 (237.6) |
|  | Skew. / kurt. | 0.78 / 3.01 | 3.58 / 19.71 | 3.09 / 13.53 | 4.21 / 24.80 | 3.80 / 21.13 |
|  | RCV<sub>*Q*</sub> (%) | 25.2 | 43.8 | 49.9 | 37.2 | 43.4 |
| *u*<sub>*c*</sub> | Median (IQR) | −11.8 (20.2) | −30.4 (67.7) | −27.4 (67.5) | −18.5 (44.9) | −20.6 (49.5) |
|  | Skew. / kurt. | −0.77 / 3.77 | −3.44 / 17.92 | 1.12 / 16.14 | −1.51 / 29.52 | −1.54 / 26.00 |
| *r*/*H* | Median (IQR) | 0.167 (0.053) | 0.273 (0.158) | 0.319 (0.221) | 0.228 (0.113) | 0.244 (0.139) |
|  | Skew. / kurt. | 0.82 / 2.82 | 3.99 / 23.91 | 3.03 / 13.19 | 4.12 / 24.05 | 3.88 / 22.26 |
|  | RCV<sub>*Q*</sub> (%) | 23.8 | 43.4 | 52.0 | 37.2 | 42.7 |
| *u*<sub>*c*</sub>/*H* | Median (IQR) | −0.007 (0.012) | −0.017 (0.038) | −0.017 (0.043) | −0.011 (0.027) | −0.012 (0.029) |
|  | Skew. / kurt. | −0.67 / 3.29 | −3.78 / 21.38 | 1.02 / 15.53 | −1.31 / 28.51 | −1.49 / 25.53 |

Height-normalised radius and centre horizontal coordinate both exhibited non-Gaussian distributions as well (shown in Figure 6c–d). Their skewness and kurtosis changed little from their non-normalised equivalents. Their median (IQR), skewness and kurtosis are given in Table 5; mean ± SD was 0.31 ± 0.25 for the radius and -0.024 ± 0.062 for the centre horizontal coordinate.

A t-location-scale distribution was fitted to each parameter, which are shown juxtaposed to each histogram in Figure 6.

Due to substantial positive skewness and kurtosis in radius distributions, relative variability is quantified using the quartile-based robust coefficient of variation (RCV*_Q_*, section 2.5.1), yielding 43.4% for the non-normalised radius and 42.7% for the height-normalised radius in the pooled sample (Table 5). Within datasets, RCV*_Q_* of the radius ranged from 25.2% (Dataset 1) to 49.9% (Dataset 3). Conventional coefficient of variation (CV) is additionally reported for comparison: 78.37% and 79.26% for the non-normalised and height-normalised radius, respectively.

### 3.4. Pairwise Anthropometric Associations

Descriptive statistics of age, height, mass and sex are given per dataset in Table 1. Leg, thigh, and shank lengths averaged 0.865 ± 0.072 m (range 0.616–1.070 m), 0.473 ± 0.044 m (range 0.358–0.590 m), and 0.396 ± 0.035 m (range 0.267–0.488 m), respectively, across the pooled sample (*N_L_* = 467 legs). Pooled pairwise associations between the circular arc parameters and the anthropometric variables were small for both the raw and the height-normalised parameters (all |*ρ*| ≤ 0.23 and *R*^2^ *<* 0.04 for the numeric variables); they are reported descriptively in Appendix C. Anthropometric effects were estimated jointly in the covariate models (section 3.5.3).

### 3.5. Mixed-Effects Modelling of the Radius

The mixed-effects model in section 2.5.2 was estimated on the 4 datasets, 246 participants out of *N_p_*= 249 and 453 observations (legs) out of *N_L_* = 467. Note that 14 observations were excluded from the model due to *R_H_ >* 1, of which 3 participants were excluded entirely, as both their legs fell above the threshold (6 of the 14 observations).

#### 3.5.1. Initial Model (Raw Radius Scale)

Initially we fitted the model to *r* (mm). The resultant model components are described in Table 6. 95% CIs and SEs are presented.

**Table 6:** Summary of mixed-effects model parameter estimation for the *r* model (mm). Fixed effects (dataset) coefficients are relative to the grand mean. Standard errors (SE) and 95% confidence intervals (CI) were estimated using a case-bootstrapping method. The CIs are presented as [lower upper].

| Parameter | Estimate | SE | CI |
|---|---|---|---|
| Grand mean | 465 | 12 | [443 489] |
| Dataset 1 effect | -157 | 18 | [-190 -121] |
| Dataset 2 effect | 80 | 22 | [39 123] |
| Dataset 3 effect | 111 | 24 | [65 161] |
| Dataset 4 effect | -35 | 17 | [-67 -2] |
| *τ*<sup>2</sup> (mm<sup>2</sup>) | 17932 | 6976 | [5357 32285] |
| *σ*<sup>2</sup> (mm<sup>2</sup>) | 30975 | 6587 | [19440 45283] |
| ICC | 0.367 | 0.120 | [0.121 0.586] |

Leaving out one participant at a time, the ICC ranged from 0.272 to 0.440. The extremes arose from two participants, each with at least one leg just below the *R_H_*cutoff, which changed the ICC by −0.094 and +0.074, respectively.

The participant random intercept was supported (LRT: *χ*^2^ = 24.2, *p <* 10^−6^; conservative, see section 2.5.4). Normality was rejected for the pooled *r*, the Pearson residuals and the random effects (Anderson-Darling: *A*^2^ = 22, 19, 6.5, respectively; all *p <* 0.001), and residual variance differed between datasets (Brown-Forsythe: *W* = 7.39, *p <* 10^−4^), with residual spread increasing with fitted value (Figure 7a).

**Figure 7:**
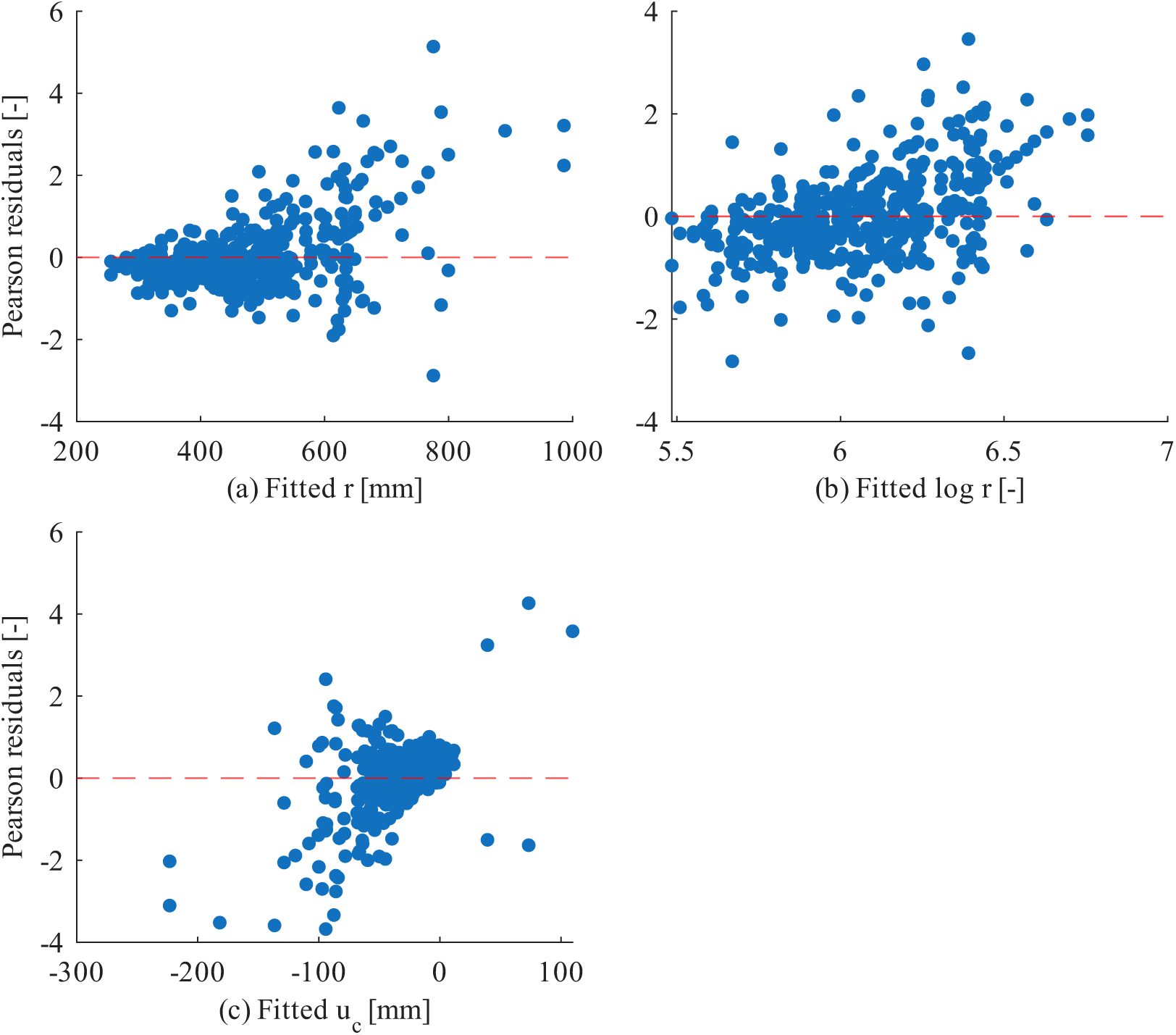
Pearson residuals versus fitted values of the mixed-effects models (dataset fixed effect, participant random intercept) of (a) the roll-over shape (ROS) radius *r* (mm), (b) log *r* (*r* in mm; fitted values and residuals on the naturallog scale, so they are not comparable in magnitude with (a)) and (c) the centre horizontal coordinate *u_c_*(mm, positive anterior to the ankle); 453 legs of 246 participants, after excluding legs with *r/H >* 1 (*H*: participant height). Dashed line: zero residual.

#### 3.5.2. Log-Transformed Model

Fitting the response as log *r*, the resultant model components are presented in Table 7. Again, a case bootstrapping method was used for parameter inference estimation.

**Table 7:** Summary of mixed-effects model parameter estimation for the log *r* model (*r* in mm). Fixed effects (dataset) coefficients are relative to the grand mean. Standard errors (SE) and 95% confidence intervals (CI) were estimated using a case-bootstrapping method. The CIs are presented as [lower upper].

| Parameter | Estimate | SE | CI |
|---|---|---|---|
| Grand mean | 6.044 | 0.024 | [5.999 6.090] |
| Dataset 1 effect | -0.340 | 0.048 | [-0.433 -0.245] |
| Dataset 2 effect | 0.173 | 0.038 | [0.100 0.248] |
| Dataset 3 effect | 0.238 | 0.041 | [0.158 0.316] |
| Dataset 4 effect | -0.071 | 0.033 | [-0.135 -0.008] |
| *τ*<sup>2</sup> | 0.066 | 0.015 | [0.036 0.094] |
| *σ*<sup>2</sup> | 0.090 | 0.014 | [0.064 0.118] |
| ICC<sub>*log*</sub> | 0.423 | 0.082 | [0.249 0.571] |
| ICC | 0.405 | 0.080 | [0.235 0.551] |

For comparison and interpretability purposes, the log scale ICC (ICC*_log_*, Table 7) was converted to the original *r* scale using the law of total variance, as shown in Equation 8. The casebootstrapped SE and 95% CI was also rescaled. These rescaled results are also presented in Table 7.

The random intercept was again supported (LRT: *χ*^2^ = 36.3, *p <* 10^−8^). Anderson-Darling statistics were lower than for the *r* model for the pooled log *r*, the Pearson residuals and the random effects (*A*^2^ = 3.5, 4.1, 0.86, respectively), but normality was still rejected (pooled log *r* and residuals: *p <* 0.001; random effects: *p* = 0.027). Heteroscedasticity was reduced but not removed (Brown-Forsythe: *W* = 4.80, *p* = 0.0027; Figure 7b).

Because heteroscedasticity remained, observation-level weights were considered (section 2.5.4). *e*_r_ increased with *n*_steps_ (Spearman *ρ* = 0.586, *p <* 10^−42^; log-log exponent 0.297, *R*^2^ = 0.34), so it reflects the number of steps in each fit rather than an independent measure of precision. The decrease in log *r* variance with *n*_steps_ (power-law exponent −0.28, *R*^2^ = 0.60) largely disappeared after subtracting dataset means (exponent −0.062, *R*^2^ = 0.09; bin-size-weighted slope −0.106, *S E* = 0.108, *p* = 0.40), so the unweighted model was retained.

Leaving out one participant at a time, ICC*_log_* ranged from 0.387 to 0.465. The extremes arose from the same two participants as in the *r* model, but with a smaller influence (−0.037 and +0.041, compared with −0.094 and +0.074).

Estimated within each dataset, ICC*_log_* was 0.82 (95% CI [0.59, 0.91]) in Dataset 1, 0.07 [0.00, 0.39] in Dataset 2, 0.35 [0.00, 0.74] in Dataset 3, and 0.55 [0.36, 0.71] in Dataset 4 (Figure 8a). The CIs of Datasets 1 and 2 did not overlap. In Datasets 2 and 3, 35% and 9% of the bootstrap resamples, respectively, estimated *τ*^2^ = 0. Dataset 3 also had the lowest proportion of participants with both legs measured (24 of 49).

**Figure 8:**
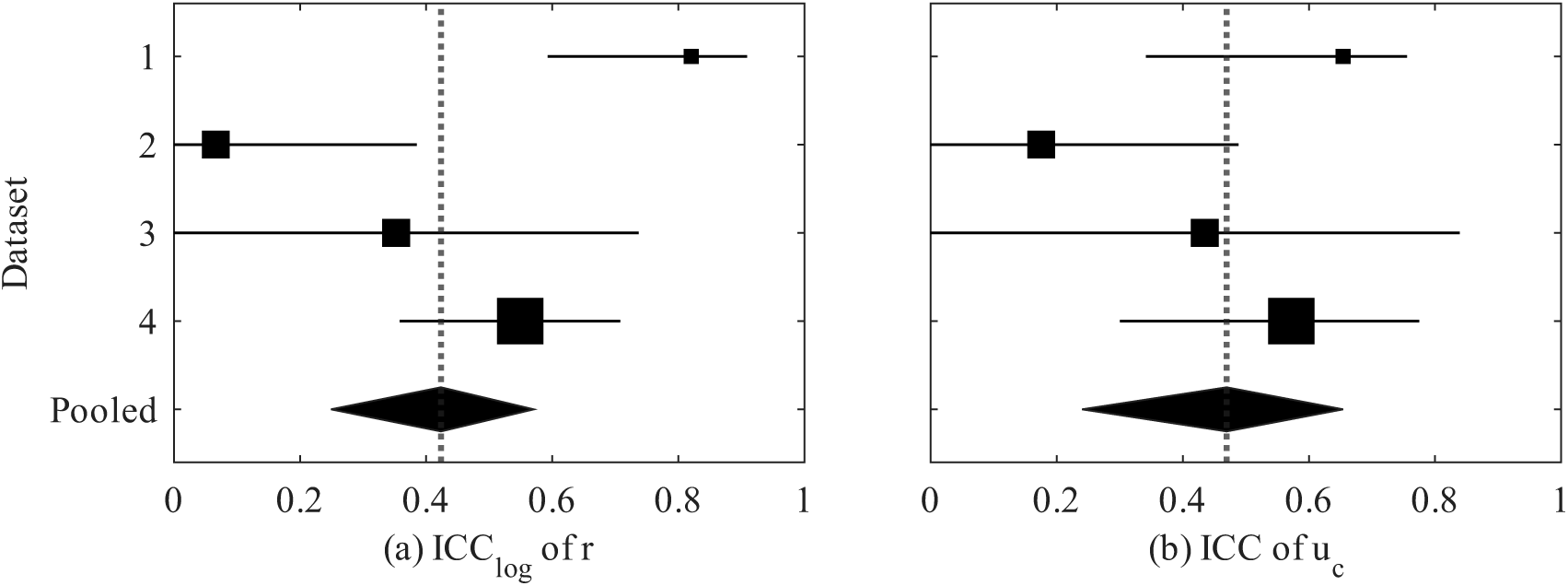
Intraclass correlation coefficient (ICC) of (a) the roll-over shape (ROS) radius *r*, from the model of log *r* (ICC*_log_*: ICC on the log scale), and (b) the centre horizontal coordinate *u_c_* (mm, positive anterior to the ankle). Squares: ICC estimated within each dataset (area proportional to the number of participants); diamond and dotted line: pooled mixed-effects model with dataset fixed effects. Horizontal bars and diamond width: 95% case-bootstrap confidence intervals (CIs; 5000 resamples, participants drawn with replacement within each dataset). Number of participants / legs per dataset: 1, 14 / 28; 2, 48 / 94; 3, 49 / 73; 4, 135 / 258; pooled, 246 / 453.

#### 3.5.3. Covariate Models

The covariate models of section 2.5.3 are summarised in Table 8. Added jointly, age, sex, log *H* and log *M* accounted for none of the between-participant variance, and ICC*_log_* was unchanged. No individual anthropometric effect could be distinguished from zero (Table F.13), and there was no evidence that the anthropometric effects differed between datasets (LRT *χ*^2^ = 13.0, *p* = 0.37).

**Table 8:**
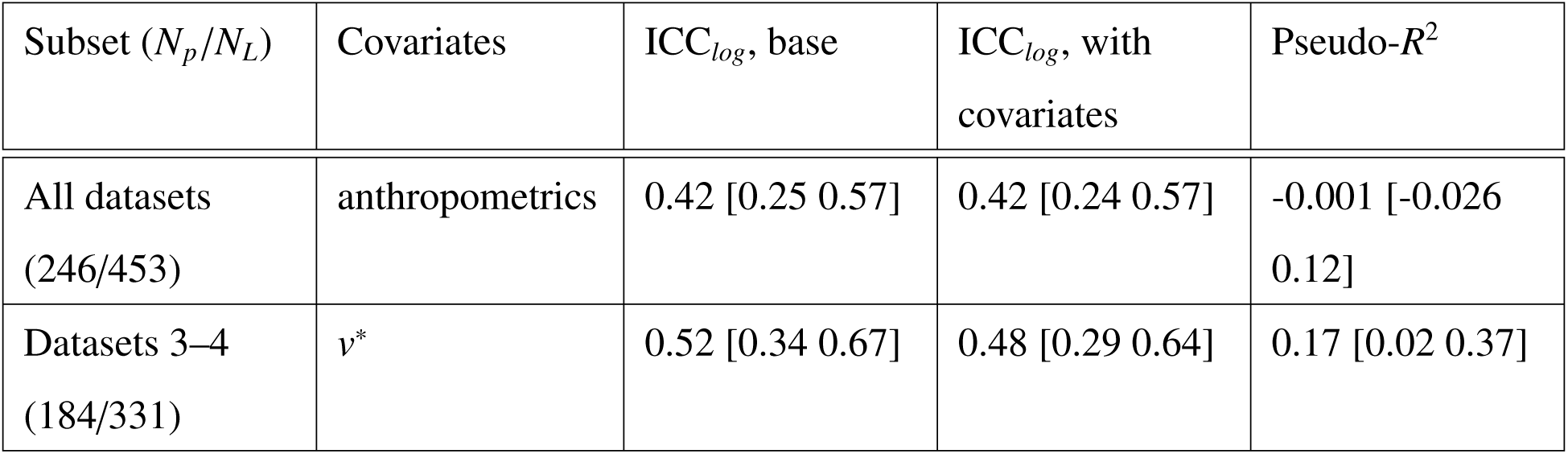
Covariate models of log *r* (section 2.5.3). Each covariate set is added to the base model (Equation 6) fitted on the same legs, and pseudo-*R*^2^ is computed against it. *N_p_/N_L_*: participants/legs. ICC*_log_* is given for the base model and for the model with the covariates. Estimates are given with 95% case-bootstrap CIs (5000 resamples) as [lower upper]. Anthropometrics: age, sex, log *H* and log *M*. *v*^∗^: dimensionless walking speed. Coefficients are given in Table F.13; sensitivity analyses in Table F.14.

| Subset (*N*<sub>*p*</sub>/*N*<sub>*L*</sub>) | Covariates | ICC<sub>*log*</sub>, base | ICC<sub>*log*</sub>, with covariates | Pseudo-*R*<sup>2</sup> |
|---|---|---|---|---|
| All datasets (246/453) | anthropometrics | 0.42 [0.25 0.57] | 0.42 [0.24 0.57] | -0.001 [-0.026 0.12] |
| Datasets 3–4 (184/331) | *v*<sup>\*</sup> | 0.52 [0.34 0.67] | 0.48 [0.29 0.64] | 0.17 [0.02 0.37] |

Proportional scaling of the radius with height was not supported: the slope of log *r* on log *H* was 0.26 (95% CI [−0.26, 0.77]), and the CI excluded 1. The result was similar without the *R_H_* ≤ 1 exclusion (0.23 [−0.37, 0.83]). Adjusted for age, sex and log *M*, the height elasticity was imprecisely estimated (Table F.13).

In the self-selected-speed datasets (3 and 4), faster walking was associated with a larger log *r* (*β* = 2.14 per unit *v*^∗^, 95% CI [1.14, 3.09]), with no evidence that the slope differed between the two datasets (LRT *p* = 0.17). Speed accounted for an imprecisely estimated share of their between-participant variance, and their ICC*_log_* decreased slightly (Table 8).

Three sensitivity and modelling checks are reported in Appendix F. There was no systematic leg-side effect, supporting the pooling of both legs without a side term. Replacing log *H* by log leg length, and fitting Dataset 4 alone, gave results similar to those of the pooled anthropometric and speed models (Table F.14).

As in the base model, the residuals of all covariate models were non-normal, and in the alldataset models the residual variance differed between datasets (Brown-Forsythe *p* = 0.003); in the Dataset 3–4 speed model it did not (*p* = 0.47). Age showed no evidence of curvature, and log *H* none in the all-dataset model. A quadratic term was supported for log *M* in the all-dataset model (95% CI of the quadratic coefficient [−0.99, −0.06]) and, in Dataset 4 alone, for log *M*, log *H* and, marginally, *v*^∗^ ([0.04, 23.4]), but not for *v*^∗^ in Datasets 3–4 ([−1.3, 17.5]); the curvature was carried by a few participants at the extremes of body size and speed, so the linear coefficients should be read as average slopes. Variance inflation factors of the covariates were at most 2.7 (log *H* 2.7, log *M* 2.5). Leaving out one participant at a time, pseudo-*R*^2^ ranged from −0.009 to 0.013 for anthropometrics, and from 0.12 to 0.19 for speed in Datasets 3–4.

### 3.6. Mixed-Effects Modelling of the Centre Horizontal Coordinate

The mixed-effects model of section 2.5.2 was fitted to *u_c_* on the same 246 participants and 453 legs as the radius models. Its components are given in Table 9. The grand mean of *u_c_* was −33.8 mm (95% CI [−40.1, −27.9]), posterior to the ankle, and between-participant differences accounted for 47% of its variance (ICC 0.469 [0.240, 0.654]). The participant random intercept was supported (LRT: *χ*^2^ = 40.1, *p <* 10^−9^). Leaving out one participant at a time, the ICC ranged from 0.371 to 0.509; the lower extreme arose from a single Dataset 4 participant.

**Table 9:** Summary of mixed-effects model parameter estimation for the *u_c_* model (mm). Fixed effects (dataset) coefficients are relative to the grand mean. Standard errors (SE) and 95% confidence intervals (CI) were estimated using a case-bootstrapping method. The CIs are presented as [lower upper].

| Parameter | Estimate | SE | CI |
|---|---|---|---|
| Grand mean | -33.8 | 3.1 | [-40.1 -27.9] |
| Dataset 1 effect | 19.2 | 4.1 | [10.8 27.2] |
| Dataset 2 effect | -12.2 | 5.4 | [-23.2 -1.9] |
| Dataset 3 effect | -13.6 | 6.6 | [-27.1 -1.3] |
| Dataset 4 effect | 6.6 | 4.5 | [-2.1 15.5] |
| *τ*<sup>2</sup> (mm<sup>2</sup>) | 1649 | 587 | [659 2878] |
| *σ*<sup>2</sup> (mm<sup>2</sup>) | 1864 | 361 | [1225 2633] |
| ICC | 0.469 | 0.109 | [0.240 0.654] |

Estimated within each dataset, the ICC was 0.65 (95% CI [0.34, 0.76]) in Dataset 1, 0.18 [0.00, 0.49] in Dataset 2, 0.43 [0.00, 0.84] in Dataset 3, and 0.57 [0.30, 0.77] in Dataset 4 (Figure 8b). In Datasets 2 and 3, 12% and 13% of the bootstrap resamples, respectively, estimated *τ*^2^ = 0. As for the radius, the lowest ICC was found in Dataset 2. Anderson-Darling tests rejected normality for the pooled *u_c_*, the Pearson residuals and the random effects (*A*^2^ = 24, 21, 7.2, respectively; all *p <* 0.001), and the residual variance differed between datasets (Brown-Forsythe: *W* = 4.51, *p* = 0.004). The residual spread increased as the fitted *u_c_*became more posterior (Figure 7c). Without the *R_H_* ≤ 1 exclusion, the 14 nearly flat arcs dominated the estimates: the ICC of Dataset 3 collapsed to zero, and leaving out a single participant changed the pseudo-*R*^2^ of speed by up to 0.68.

The covariate models of *u_c_* are summarised in Table 10. Anthropometrics accounted for none of the between-participant variance, and no individual effect could be distinguished from zero (Table F.13). Proportional scaling at the sample’s mean ratio of *u_c_* to height would correspond to a height slope of about −20 mm per m, inside the CI of the estimated slope. There was no evidence that the anthropometric effects differed between datasets (LRT *χ*^2^ = 10.0, *p* = 0.61).

**Table 10:**
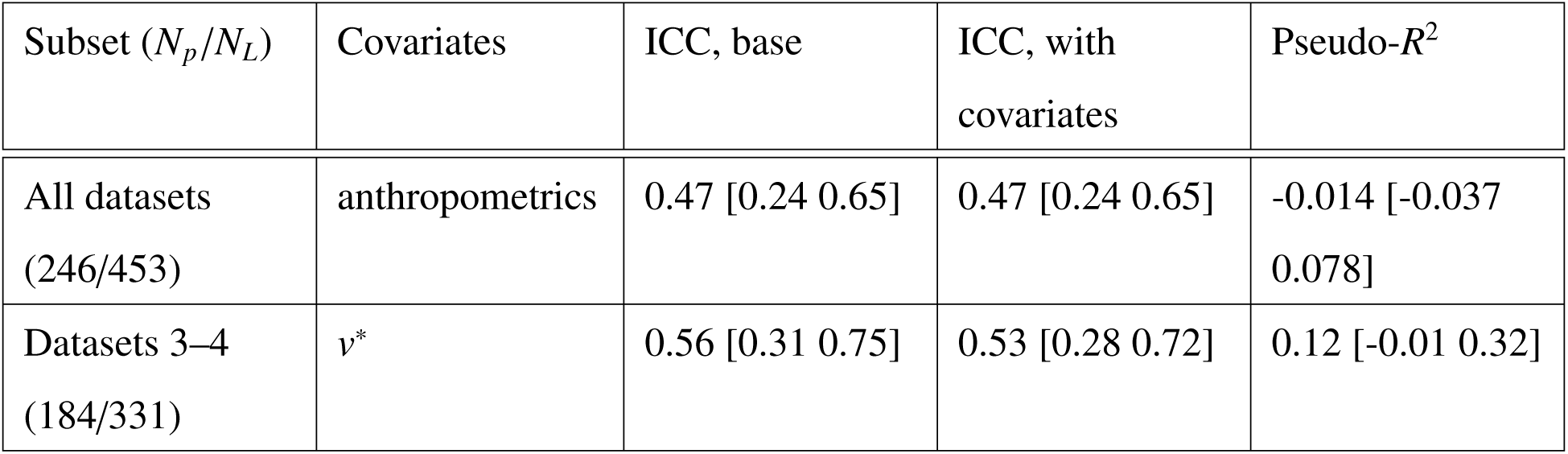
Covariate models of *u_c_* (section 2.5.3). Each covariate set is added to the base model (Equation 6) fitted on the same legs, and pseudo-*R*^2^ is computed against it. *N_p_/N_L_*: participants/legs. The ICC is given for the base model and for the model with the covariates. Estimates are given with 95% case-bootstrap CIs (5000 resamples) as [lower upper]. Anthropometrics: age, sex, height and mass. *v*^∗^: dimensionless walking speed. Coefficients are given in Table F.13; sensitivity analyses in Table F.15.

| Subset (*N<sub>p</sub>*/*N<sub>L</sub>*) | Covariates | ICC, base | ICC, with covariates | Pseudo-*R*<sup>2</sup> |
|---|---|---|---|---|
| All datasets (246/453) | anthropometrics | 0.47 [0.24 0.65] | 0.47 [0.24 0.65] | -0.014 [-0.037 0.078] |
| Datasets 3–4 (184/331) | *v*<sup>\*</sup> | 0.56 [0.31 0.75] | 0.53 [0.28 0.72] | 0.12 [-0.01 0.32] |

In the self-selected-speed datasets (3 and 4), faster walking was associated with a more posterior centre (*β* = −260 mm per unit *v*^∗^, 95% CI [−455, −78]), with no evidence that the slope differed between the two datasets (LRT *p* = 0.25). Speed accounted for an imprecisely estimated share of their between-participant variance, and their ICC decreased slightly (Table 10).

The right centre was slightly more posterior than the left, a difference not reproduced in Dataset 4 alone (Table F.13). Replacing height by leg length, and fitting Dataset 4 alone, gave results similar to those of the pooled models (Table F.15).

The residuals of all *u_c_* covariate models were non-normal, and in the all-dataset model the residual variance differed between datasets (Brown-Forsythe *p* = 0.003); in the Dataset 3–4 speed model it did not (*p* = 0.19). Height and mass showed no evidence of curvature. A quadratic term was supported for *v*^∗^ in both speed models, with the relation steepening at the fastest speeds, where few legs were available, so the linear speed coefficients should be read as average slopes; a weaker curvature in age was carried by the participants youngest relative to their dataset mean. Variance inflation factors were at most 2.6. Leaving out one participant at a time, pseudo-*R*^2^ ranged from −0.027 to −0.005 for anthropometrics, and from 0.06 to 0.15 for speed in Datasets 3–4.

## 4. Discussion

### 4.1. ROS Parameter Distributions

Because all retained steps of a leg were pooled into a single point cloud, *e*_r_ reflects both the deviation of each step from a circular arc (intra-step variability) and the step-to-step variation of the ROS (inter-step variability), and therefore should not be read as a pure measure of fit quality; this is consistent with its positive association with the number of steps per leg. The non-normality observed in *e*_r_, particularly pronounced in Dataset 4, likely reflects a combination of inter-individual variability, inter-dataset differences in measurement protocol, and a subset of lower-quality steps rather than a single dominant cause; disentangling these contributions is left to future work.

The radius and centre horizontal coordinate both exhibited heavy-tailed, non-Gaussian distributions, with high kurtosis and skewness. A t-location-scale distribution provided a visually better qualitative fit than a normal distribution for both parameters, suggesting that ROS metrics are inherently prone to outliers rather than approximately Gaussian, and that summary statistics relying on normality (e.g., mean ± SD) may understate the true spread of these measures. The RCV*_Q_* of the radius, which is more robust to outliers than the conventional CV, nonetheless remained substantial (∼ 43%), indicating that high relative variability is not solely an artefact of a small number of extreme values but a genuine feature of the underlying distribution. The negative skewness of the centre horizontal coordinate reflects a majority of centre horizontal coordinate values lying posterior to the ankle in our dataset. The mixed-effects model confirms this beyond the skewness of the pooled distribution: after accounting for dataset and for the correlation between legs, the mean *u_c_* was −33.8 mm (95% CI [−40.1, −27.9]; section 3.6). This contrasts with prior reports of the knee-ankle-foot ROS, computed in a hip-ankle-based CS, which found small positive median height-normalised values (0.006–0.011) [1, 21], corresponding to a predominantly anterior position. This discrepancy may partly reflect the difference between the hip-ankle axis defining the knee-ankle-foot ROS and the knee-ankle (shank) axis defining the ankle-foot ROS used here: since the hip and knee are not generally collinear with the ankle during stance, particularly under stance-phase knee flexion, the two coordinate definitions need not agree on the anteroposterior position of the roll-over centre even when applied to similar gait patterns.

### 4.2. Height Normalisation and Anthropometric Scaling

Normalising radius by body height did not meaningfully reduce the variability of the radius, nor did it strengthen the pooled associations with anthropometric variables (all *R*^2^ *<* 0.04 for both the raw and the height-normalised parameters; Appendix C), and the normalised radius retained a distribution comparable to the raw parameter (skewness 3.88 vs 3.80; RCV*_Q_* 42.7% vs 43.4%). This suggests that height-normalisation, at least for this parameter, may not isolate a scale-invariant quantity so much as divide by a comparatively low-variance constant: participant height was modest in variability within each dataset (CV ≈ 5 − 12.5%, computed from Table 1, prior to step-level screening) relative to the substantially larger variability observed in radius and height-normalised radius through the RCV*_Q_*values, and CV = 78.37% and 79.26%, respectively. Because height varies far less than radius, dividing by height can change the relative variability of the radius by at most roughly the CV of height, and only to the extent that radius and height are correlated. The unadjusted scaling analysis is consistent with this: the slope of log *r* on log *H* was 0.26 (95% CI [−0.26, 0.77]), which excludes proportional scaling. Dividing by height may therefore over-correct, leaving the height-normalised radius negatively dependent on height and on correlated measures such as leg and shank length: the pooled rank correlation with height was 0.03 for the raw radius but −0.14 for the height-normalised radius (Table C.11). The same applies to sex: the sex difference was negligible for the raw radius (*r_rb_* = −0.05) but small for the height-normalised radius (*r_rb_* = −0.17), consistent with the division by height, which was lower on average in women (1.61 vs 1.75 m). These pooled analyses treat legs as independent and are reported descriptively; adjusted for each other, for dataset, and for the correlation between legs, no individual anthropometric effect could be distinguished from zero (section 4.3). This challenges an implicit assumption in prior literature that height-normalisation meaningfully accounts for anthropometric scaling of ROS radius. Because the height elasticity adjusted for age, sex and mass was imprecisely estimated (0.78 [−0.18, 1.76]), the data do not establish how the radius scales with body size, only that proportional scaling with height was not supported.

### 4.3. Inter-Participant Variability of the Radius

#### 4.3.1. Rationale for the Log Transformation

The initial model fit on raw *r* showed a clear funnel pattern in the residual-vs-fitted plot (Figure 7a), visibly increasing across the range of fitted values. This pattern is the classic signature of multiplicative, rather than additive, measurement error: variance that scales with the mean of the response rather than remaining constant. A log transform is the standard remedy for this error structure, and refitting the model on log *r* substantially reduced both the residual heteroscedasticity (Brown-Forsythe *W* dropping from 7.39 to 4.80, with the residual-vs-fitted plot showing a visibly attenuated funnel; Figure 7b) and the non-normality of the pooled response, Pearson residuals, and random effects (AD *A*^2^ statistics dropping from 22 to 3.5 for the pooled response, and similarly 19 and 6.5 to 4.1 and 0.86 for the residuals and random effects, respectively).

The log transformation also reduced the influence of individual participants on the ICC: the largest change when leaving out one participant was 0.094 for *r* but 0.041 for log *r*, caused in both models by the same two participants. Because the fitted radius scales approximately inversely with arc span, legs with nearly flat ROSs just below the cutoff are stretched on the raw scale, and the log transformation compresses them.

It is worth emphasising that this improvement was substantial but not complete: the Brown-Forsythe test still rejected homoscedasticity for the log-transformed model, and the AD test still rejected normality of the residuals (*A*^2^ = 4.1, *p <* 0.001) and, marginally, of the random effects (*A*^2^ = 0.86, *p* = 0.027). Under such distributional misspecification, the fixed-effect estimates remain virtually unbiased [34], and model-based standard errors and tests were replaced with casebootstrapped inference [34]. The covariance structure, however, is not fully correctly specified: the residual variance differed between datasets, whereas the model assumes a common one. The pooled ICC should therefore be read as a summary averaged over datasets. The dataset-specific ICCs (Figure 8a) differed substantially, from 0.07 in Dataset 2 to 0.82 in Dataset 1.

#### 4.3.2. Interpretation of the Weighting Investigation

Because the residual variance appeared to scale with the fitted value even after transformation, we considered whether an observation-level weighting scheme, using the inverse variance as weights [35], could further address the remaining heteroscedasticity.

The number of steps used to fit each circular arc (*n*_steps_) was a natural candidate, under the hypothesis that arcs fit from more steps yield more precise radius estimates. However, *n*_steps_ varies enormously between datasets (Dataset 1: 191.6 ± 28.2 vs. Dataset 4: 4.8 ± 2.0 steps per participant), raising the concern that any apparent relationship between *n*_steps_ and residual variance might simply reflect between-dataset differences already captured by the dataset ID fixed effect, rather than a genuine within-dataset precision effect. Testing this directly, the unadjusted power-law exponent relating log *r* variance to *n*_steps_ (−0.28, *R*^2^ = 0.60) collapsed to −0.062 (*R*^2^ = 0.09), once dataset means were subtracted out. Also, a bin-size-weighted slope was indistinguishable from zero (−0.106, SE 0.108, *p* = 0.40). This indicates that the apparent precision gain from additional steps is largely, if not entirely, an artefact of confounding with dataset ID, and that *n*_steps_ does not carry meaningful additional information about measurement precision once dataset is accounted for.

*e*_r_ was also considered but rejected, since it increased with *n*_steps_ (Spearman *ρ* = 0.586) and thus reflects pooled inter-step variability rather than independent precision (section 4.1).

#### 4.3.3. Robustness of the Inter-Participant Variance Estimate

The central finding of this analysis is that a substantial and statistically robust proportion of variance in log *r* is attributable to between-participant differences. Several convergent lines of evidence support this conclusion. First, the case-bootstrapped confidence interval for the log-scale ICC ([0.249, 0.571]), which does not rely on the normality assumptions that were not fully met, excludes zero comfortably and is not close to the parameter boundary, supporting a stable, wellidentified estimate for our primary model specification. Second, the LRT comparing models with and without the random participant intercept was decisively significant for both the raw and logtransformed models (*χ*^2^ = 24.2, 36.3, respectively, both *p <* 10^−6^). Because the null hypothesis in this test (*τ*^2^ = 0) lies on the boundary of the parameter space, the standard *χ*^2^ reference distribution is conservative [33], so a boundary correction would only strengthen this result; however, the test’s reference distribution also assumes normally distributed random effects and residuals [34], so we regard it as supporting rather than primary evidence. Third, within the largest dataset (Dataset 4, 135 participants), which avoids cross-dataset pooling and the *n*_steps_-related confound discussed above, ICC*_log_* = 0.55 (95% CI [0.36, 0.71]), compatible with the pooled estimate. The datasetspecific ICCs were, however, heterogeneous (Figure 8a). They did not follow the speed protocol: the highest ICC was found in a metronome-cued dataset (Dataset 1) and the lowest in another (Dataset 2). The wide CIs of Datasets 2 and 3 mean that these estimates are imprecise rather than evidence of an absent participant effect. Fourth, anthropometrics accounted for none of the between-participant variance (pseudo-*R*^2^ −0.1%, 95% CI −3 to 12%), and in Datasets 3–4 selfselected walking speed accounted for an imprecisely estimated minority of it (17%, 2–37%); after adjustment for anthropometrics, ICC*_log_* remained 0.42 (95% CI [0.24, 0.57]), and 0.48 [0.29, 0.64] after adjustment for speed in the self-selected-speed datasets.

Taken together, we interpret this as evidence that a substantial share of the variance in log *r* is shared between the two legs of a participant within a session: after accounting for dataset, between-participant differences accounted for 42% of the variance in log *r*, with the remainder reflecting within-participant (left-right) differences and arc-fit uncertainty, which the present design cannot separate, and also notwithstanding residual non-normality and heteroscedasticity discussed above. Self-selected speed was associated with part of this between-participant variance; whether the remainder reflects a stable individual trait or other session-level factors requires repeatedsession data.

#### 4.3.4. Dataset Fixed Effects

Dataset ID was retained as a fixed, rather than random, effect, since only four dataset levels are available, well below the number generally considered necessary for a reliable random-effects variance estimate [36]. This choice reflects our primary interest in the level-2 (participant) variance component, but it also means the reported dataset-specific coefficients describe differences among these four specific datasets and their associated protocols, rather than characterising a hypothetical population of datasets or measurement protocols more broadly. The substantial and significant differences between datasets (e.g., Dataset 1 showing a markedly lower log *r* relative to the grand mean and Dataset 3 markedly higher) may reflect genuine protocol-level differences, such as walking surface, instructed gait speed, or marker placement conventions. Footwear is one such factor that cannot be separated: only Dataset 1 was shod, so the pooled shod-versus-barefoot difference in radius (Appendix C) is equivalent to comparing Dataset 1 with the remaining datasets, and coincides with its largest negative dataset effect (−0.340 on the log scale; Table 7). However, disentangling which specific protocol factors drive these differences would require data not available in the present harmonised dataset.

#### 4.3.5. Interpretation of the Back-Transformed ICC

Because log *r* was substantially closer to normality than *r* at both the participant and residual level (AD *A*^2^ = 19 → 4.1 and 6.5 → 0.86), we report the ICC primarily on the log scale. For readers more accustomed to interpreting variability in the original units of *r*, we additionally provide a back-transformed estimate of 0.405 using the standard result for a log-normal random-intercept model. This transformation is exact only under the assumption that *r* is conditionally log-normal; given the residual non-normality documented above, this should be treated as an approximate reference value rather than an independently and exactly estimated quantity. Its close agreement with the log-scale ICC (0.423) is reassuring, though this proximity is partly a mathematical consequence of the variances involved being modest in magnitude, and should not be taken as validating the log-normality assumption itself.

### 4.4. Inter-Participant Variability of the Centre Horizontal Coordinate

The centre horizontal coordinate showed a between-participant share of variance similar to that of the radius (ICC 0.47, 95% CI [0.24, 0.65], compared with a back-transformed ICC of 0.41, 95% CI [0.24, 0.55], for the radius on its original scale), with the same heterogeneity between datasets and the lowest ICC again in Dataset 2. As for the radius, anthropometrics accounted for none of this variance (pseudo-*R*^2^ −0.014, 95% CI [−0.037, 0.078]). No individual effect, including that of height, could be distinguished from zero. Within the sample, the fore-aft position of the rollover centre therefore could not be predicted from body size or demographics. For device design, where the centre position must be specified [17, 16], this suggests that a population value scaled by body size is unlikely to capture an individual’s centre position. Whether the individual deviation is functionally relevant, and stable enough to warrant participant-specific measurement or tuning, remains to be tested.

In the self-selected-speed datasets, faster walking was associated with a more posterior centre: across the central 90% of *v*^∗^ (−0.08 to +0.10 around each dataset mean), the estimated slope corresponds to a difference of about 47 mm, larger than any of the dataset effects. Together with the larger radius at faster speeds, this indicates that the whole arc, and not only its curvature, changes with speed. The relation was not linear, steepening at the fastest speeds where data were sparse, and speed accounted for an imprecisely estimated share of the between-participant variance (12%, 95% CI −1 to 32%).

The small side difference (right − left −8.3 mm, 95% CI [−16.1, −0.7]) was not reproduced in Dataset 4 alone, and its CI nearly reaches zero. Because *u_c_* depends on the orientation of the shank axis, an asymmetry in marker placement could produce a difference of this size, so we do not interpret it as a functional asymmetry.

### 4.5. Limitations

While *PROST*supports analysis using medial markers to better estimate the centreline of the medial-lateral joint locations, two of the four datasets used here [25, 26] did not include medial marker measurements. To ensure consistency across datasets, we therefore restricted the present analysis to lateral marker trajectories throughout. Future studies using our toolbox on datasets with complete medial marker data could evaluate whether medial or mean marker trajectories yield materially different ROS estimates.

For nearly flat ROSs, the fitted radius is weakly constrained: the objective function varies little over a wide range of radii, so individual large radii (approaching or exceeding body height) are uncertain. Population summaries were therefore based on robust statistics (median and IQR).

Anthropometric effects were estimated jointly, and speed separately, within the mixed-effects model, but kinematic and kinetic predictors were not included. The between-participant variance itself was estimated with limited precision (*τ*^2^ 95% CI 0.036–0.094 on the log scale), so the shares explained by covariates are imprecise. Speed could be analysed only in the two self-selected-speed datasets, and its effect, like that of mass, may be non-linear at the extremes of the sample.

The centre horizontal coordinate depends on the orientation of the shank axis, which the radius does not: with the centre a median of 332 mm above the ankle, a 1^◦^ tilt of the axis shifts *u_c_* by about 6 mm. The lateral-only markers and the estimated knee markers used in some datasets can each tilt the axis by several degrees, so absolute values of *u_c_*, and differences between datasets in particular, are conditional on these marker conventions. Its residual spread also increased for more posterior centres, consistent with less precisely located centres in flatter arcs.

For the mixed-effects analysis specifically, several additional limitations should be noted. First, cluster sizes at the participant level were small (1–2 observations per participant for most participants), even though the number of participants (246) was large. Following McNeish et al. [36], this regime is expected to yield reasonably reliable point estimates for a random-intercept-only model, but reduced precision (wider confidence intervals) for the variance-component estimate, and would likely preclude reliably fitting a more complex random-effects structure (e.g., random slopes) had one been of interest. Second, residual non-normality and heteroscedasticity, while substantially reduced by the log transformation, were not fully eliminated; we addressed this through case-bootstrapped inference rather than pursuing more complex heteroscedastic variance structures, which were judged to offer diminishing returns relative to the additional modelling effort required. Lastly, the small number of dataset levels (four) precluded modelling dataset ID as a random effect, limiting the generalisability of the fixed dataset-level effects, as discussed above. Future work extending this analysis to a larger number of harmonised datasets could revisit both of these constraints, treating dataset as a random effect and potentially recovering additional precision in the participant-level variance estimate.

### 4.6. Summary of Findings

Across the pooled datasets, ROS parameters were well approximated by a circular arc model but exhibited substantial inter-individual variability. The distributions of both radius and centre horizontal coordinate were non-Gaussian, and anthropometric variables, analysed jointly in the mixed-effects model, accounted for none of the between-participant variance in either the radius or the centre horizontal coordinate. Normalisation by height did not meaningfully reduce variability, and proportional scaling of the radius with height was not supported, challenging common assumptions regarding ROS scaling. In the self-selected-speed datasets, faster walking was associated with a larger radius. Mixed-effects modelling of the radius confirmed that, after accounting for dataset, a substantial and robust proportion of this variability (ICC*_log_* = 0.42, back-transformed ICC = 0.41) reflects differences consistent between the two legs of a participant, a conclusion supported by case-bootstrapped confidence intervals, a leave-one-participant-out analysis and a single-dataset subset analysis, and consistent with likelihood ratio testing. The centre horizontal coordinate lay posterior to the ankle on average, showed a similar between-participant share of variance (ICC = 0.47) that anthropometrics did not explain, and moved posteriorly with faster self-selected walking. Because recording sessions were not modelled, these data do not establish whether an individual’s ROS is stable across days.

## Appendix A. Estimating the Opposite Initial Contact Time

Dataset 3 [25] did not include OIC event times. Each trial in this dataset spans a full stride from IC to IC of the same foot, encompassing the marker trajectory of both feet throughout the stride. The OIC event was therefore estimated from the heel marker height of the opposite foot.

For each trial, the heel height at IC was extracted. For each participant, the mean IC heel height was computed separately for the left and right feet. For a given trial of one foot, the mean IC heel height of the opposite foot, offset by 25 mm to ensure the threshold was reached in all trials, was used as a threshold. The OIC time was then estimated as the time at which the opposite foot’s heel marker descended below this threshold. Since force plate data spans the stance phase only, terminating at TO, the vertical GRF trace serves as a visual reference for the TO event.

As an example, Figure A.9 presents the heel height and vertical GRF of both left and right foot trials of participant 6 in Dataset 3 [25]. The solid black line shows the threshold used for OIC estimation, and the dashed lines indicate the IC, OIC, and TO event times.

**Figure A.9:**
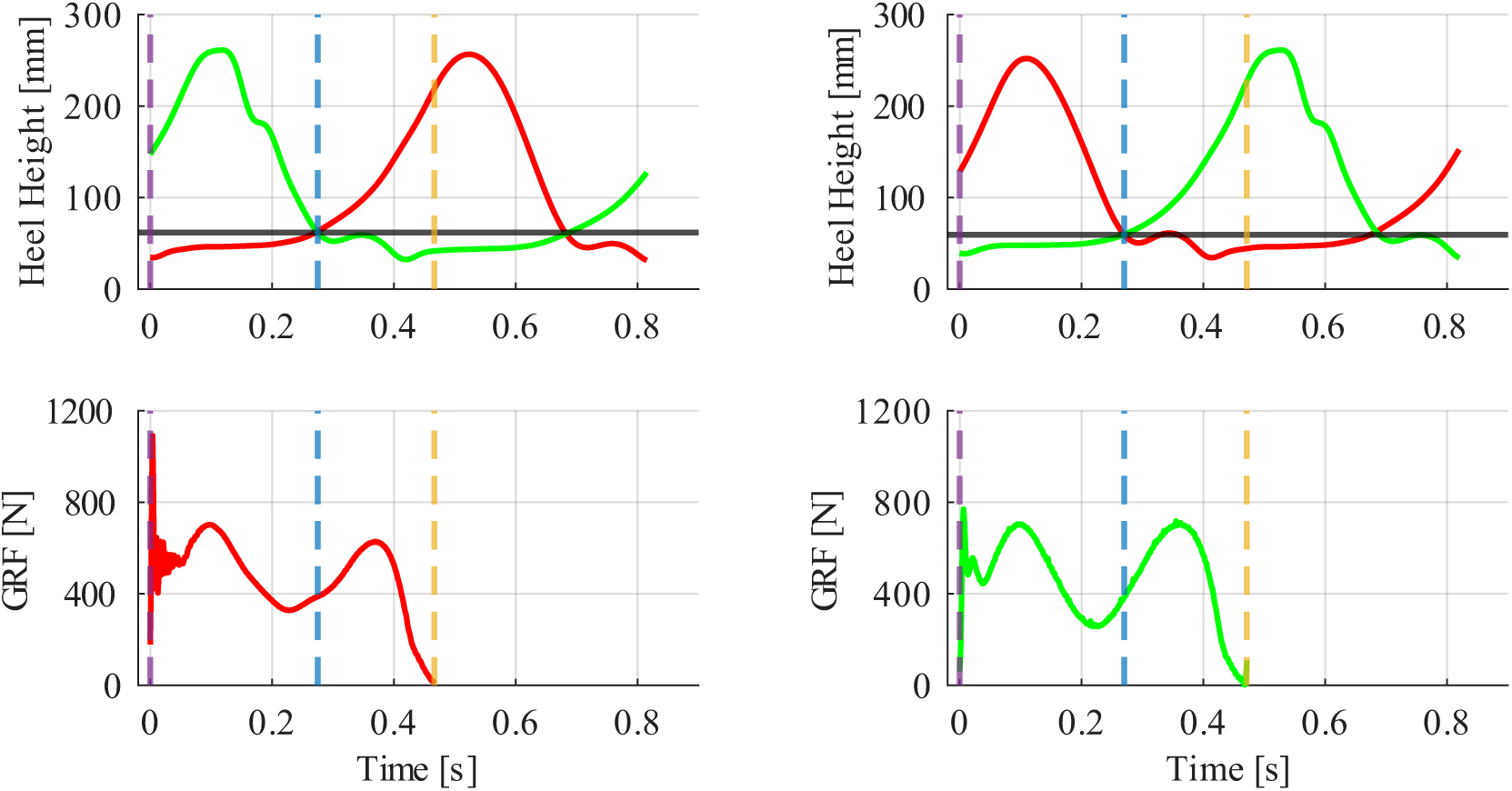
Heel height (top) and vertical ground reaction force (GRF; bottom) for a left-foot trial (left column) and a right-foot trial (right column) of participant 6 in Dataset 3 [25]. Red and green traces correspond to the left and right foot, respectively. The solid black horizontal line indicates the OIC threshold. Vertical dashed lines indicate initial contact (IC; purple), estimated opposite IC (OIC; blue), and toe-off (TO; amber).

## Appendix B. Virtual Knee Marker Estimation

The shank-based coordinate system described in subsection 2.4.1 requires knee marker trajectories, which were unavailable in datasets 2 and 3 [24, 25]. Virtual knee markers were therefore estimated from available superior and inferior markers.

Let 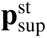 and 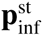 denote the positions of the superior and inferior markers relative to the knee during the standing trial. The inter-marker distance was computed as,

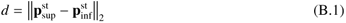

For the virtual lateral knee marker, the lateral femoral condyle and fibula head markers served as the superior and inferior references, respectively. For the virtual medial knee marker (when medial markers were present), the medial femoral condyle and tibial tuberosity were used.

The virtual knee marker trajectory during dynamic trials was then estimated as,

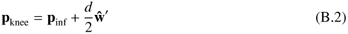

where **ŵ** ^′^ is the unit vector from the ankle to the inferior marker. A schematic of this coordinate system is shown in Figure B.10.

## Appendix C. Pairwise Associations with Anthropometric Parameters

This appendix reports the pooled pairwise associations between the circular arc parameters and the anthropometric variables summarised in section 3.4. Associations with the numeric anthropometric variables (age, height, mass, and leg, thigh, and shank lengths) were quantified using Spearman’s rank correlation coefficient *ρ* and the coefficient of determination *R*^2^ from a leastsquares linear fit (the squared Pearson correlation). Differences based on categorical parameters (that is, sex and footwear condition) were tested using the Wilcoxon rank-sum test, equivalent to the Mann–Whitney *U* test, and effect size was quantified using the rank-biserial correlation *r_rb_* [37], where a positive *r_rb_* indicates larger values in the first category (male; shod). These analyses were performed on the pooled legs, treating each leg as an independent observation. Because the two legs of a participant are correlated and the datasets differ systematically in both anthropometrics and ROS parameters, the p-values are nominal (unadjusted and likely anti-conservative) and are interpreted descriptively. Effect sizes were classified as small, moderate, or large using Cohen’s thresholds of |*ρ*| = 0.1, 0.3, and 0.5 [38], and Vargha and Delaney’s thresholds of *A* = 0.56, 0.64, and 0.71 [39], which correspond to |*r_rb_*| = 0.12, 0.28, and 0.42 through *r_rb_* = 2*A* − 1. Anthropometric effects adjusted for each other, for dataset, and for the correlation between the two legs of a participant are given by the covariate models (section 3.5.3).

**Figure B.10:**
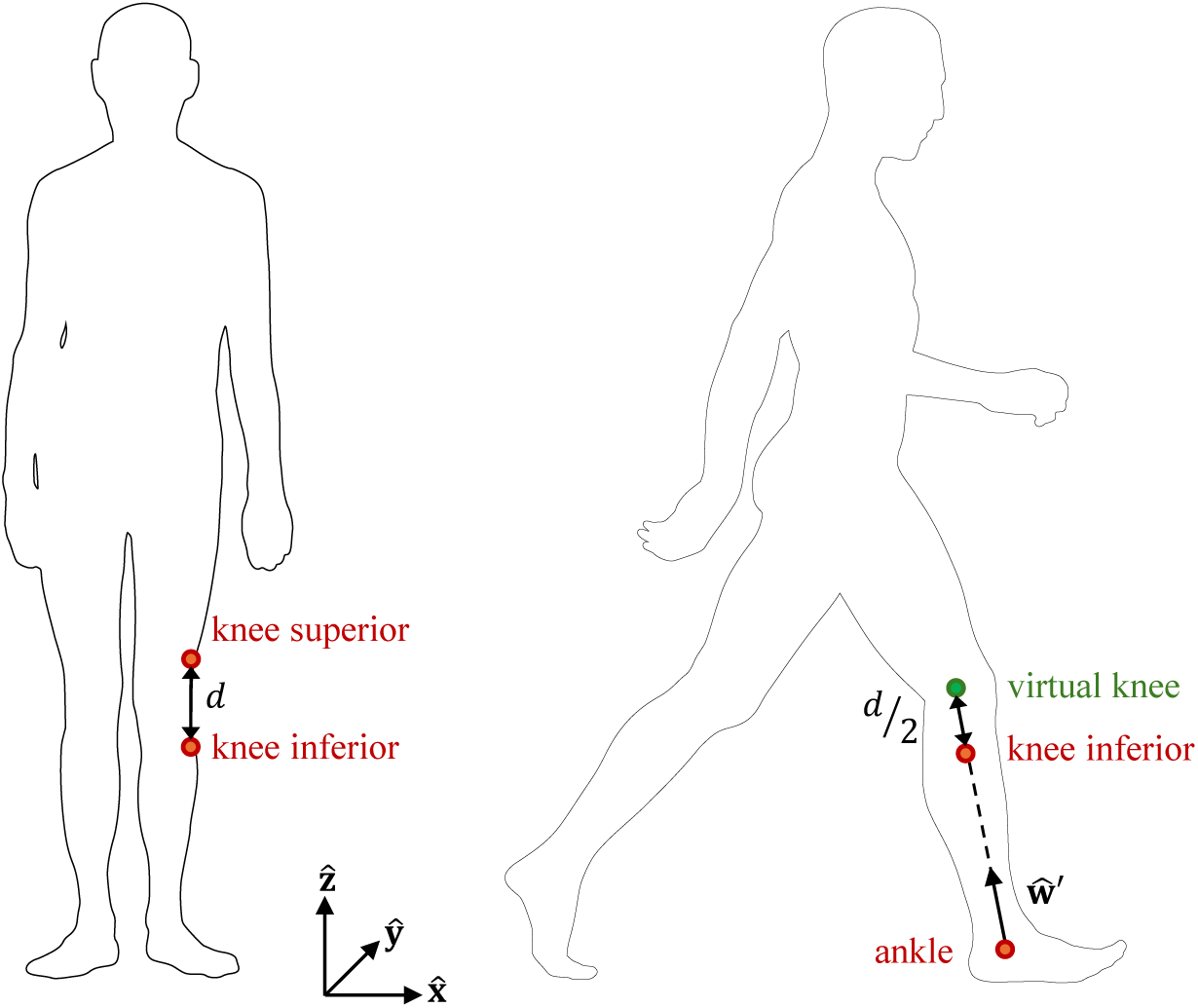
Schematic of the coordinate system (CS) used for virtual knee marker estimation. The inferior knee marker and ankle marker define the direction **ŵ** ^′^, along which half the inter-marker distance *d* is added to estimate the virtual knee marker position **p**_knee_. **x̂**, **ŷ**, **ẑ** are the axes of the laboratory CS. Figure overlays adapted from “Silhouette of a standing man” and “Silhouette of a walking man” by T.M.M. Aumala, Wikimedia Commons, 2021 (CC0).

The associations are reported primarily for the raw radius and centre horizontal coordinate. Correlating a ratio such as *r/H* with *H*, or with variables that correlate with *H* such as leg and shank length, induces a negative correlation even when *r* itself is unrelated to *H* (a ratio artefact), so associations of the height-normalised parameters with height-related variables cannot be read as associations of the ROS itself. The height-normalised results are given as well for comparison with previous studies, which reported height-normalised values [19, 1, 21, 20]. All results are summarised in Table C.11 and shown in Figure C.11–Figure C.14.

Linear fits of the raw parameters on the numeric anthropometric variables all had *R*^2^ *<* 0.02. The raw radius had a small negative correlation with shank length (*ρ* = −0.14, *p* = 0.003) and negligible correlations (|*ρ*| *<* 0.1) with age, height, mass, leg length, and thigh length. The raw centre horizontal coordinate had small positive correlations with leg length (*ρ* = 0.16, *p <* 0.001), thigh length (*ρ* = 0.11, *p* = 0.021), and shank length (*ρ* = 0.21, *p <* 10^−5^), and correlations of at most 0.10 with age, height, and mass. The sex difference was negligible for the raw radius (*z* = −0.96, *p* = 0.34, *r_rb_* = −0.05) and small for the raw centre horizontal coordinate (*z* = 2.29, *p* = 0.022, *r_rb_* = 0.12). Footwear condition differed for the raw radius (*z* = −5.16, *p <* 10^−6^, *r_rb_* = −0.58) but not clearly for the raw centre horizontal coordinate (*z* = 1.81, *p* = 0.070, *r_rb_* = 0.20).

Linear fits of the height-normalised parameters on the numeric anthropometric variables all had *R*^2^ *<* 0.04. The height-normalised radius had small negative correlations with height (*ρ* = −0.14, *p* = 0.002), mass (*ρ* = −0.18, *p <* 0.001), leg length (*ρ* = −0.12, *p* = 0.011), and shank length (*ρ* = −0.22, *p <* 10^−5^). The height-normalised centre horizontal coordinate had small positive correlations with mass (*ρ* = 0.12, *p* = 0.011) and segment lengths (leg: *ρ* = 0.18, *p <* 0.001; thigh: *ρ* = 0.12, *p* = 0.011; shank: *ρ* = 0.23, *p <* 10^−6^), and negligible correlations with height (*ρ* = 0.09, *p* = 0.044) and age (*ρ* = 0.09, *p* = 0.042). Both height-normalised parameters showed small sex differences (radius: *z* = −3.11, *p* = 0.002, *r_rb_* = −0.17; centre horizontal coordinate: *z* = 2.69, *p* = 0.007, *r_rb_* = 0.14). Footwear condition differed for the radius only (radius: *z* = −5.28, *p <* 10^−6^, *r_rb_* = −0.59; centre horizontal coordinate: *z* = 1.81, *p* = 0.071, *r_rb_* = 0.20).

**Table C.11:**
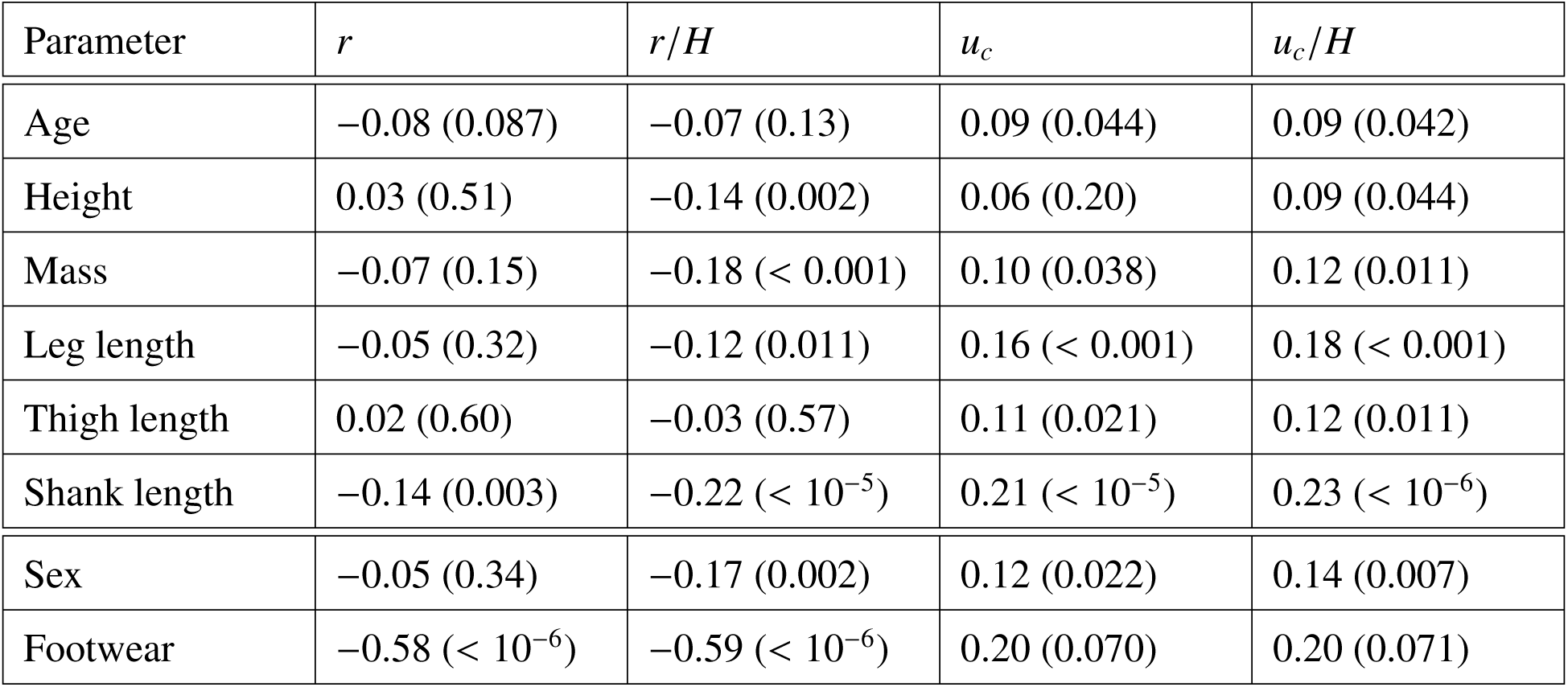
Pooled pairwise associations between the circular arc parameters and the anthropometric parameters (*N_L_* = 467 legs). Numeric parameters: Spearman *ρ* (*p*). Categorical parameters: rank-biserial correlation *r_rb_*(*p*) of the Wilcoxon rank-sum test, positive when the first category (male; shod) has larger values. *p*-values are nominal and treat legs as independent. *r*: radius; *u_c_*: centre horizontal coordinate; *H*: participant height.

| Parameter | *r* | *r*/*H* | *u<sub>c</sub>* | *u<sub>c</sub>*/*H* |
|---|---|---|---|---|
| Age | −0.08 (0.087) | −0.07 (0.13) | 0.09 (0.044) | 0.09 (0.042) |
| Height | 0.03 (0.51) | −0.14 (0.002) | 0.06 (0.20) | 0.09 (0.044) |
| Mass | −0.07 (0.15) | −0.18 (< 0.001) | 0.10 (0.038) | 0.12 (0.011) |
| Leg length | −0.05 (0.32) | −0.12 (0.011) | 0.16 (< 0.001) | 0.18 (< 0.001) |
| Thigh length | 0.02 (0.60) | −0.03 (0.57) | 0.11 (0.021) | 0.12 (0.011) |
| Shank length | −0.14 (0.003) | −0.22 (< 10<sup>−5</sup>) | 0.21 (< 10<sup>−5</sup>) | 0.23 (< 10<sup>−6</sup>) |
| Sex | −0.05 (0.34) | −0.17 (0.002) | 0.12 (0.022) | 0.14 (0.007) |
| Footwear | −0.58 (< 10<sup>−6</sup>) | −0.59 (< 10<sup>−6</sup>) | 0.20 (0.070) | 0.20 (0.071) |

Compared with the raw radius, height-normalisation introduced negative correlations with height (*ρ* = 0.03 raw vs −0.14 normalised), mass (−0.07 vs −0.18), and leg length (−0.05 vs −0.12), and the sex difference increased from negligible to small (*r_rb_* = −0.05 vs −0.17), consistent with the ratio artefact. The footwear effect on the radius was large on both scales; however, footwear condition is fully confounded with dataset ID (only Dataset 1 was shod; Table 1).

## Appendix D. Measured Walking Speed

The speed conditions in Table 1 are those reported by the source studies. They were either metronome-cued (Dataset 1 and conditions C1–C3 of Dataset 2) or self-selected, so the speed actually walked could differ from the nominal condition. Figure D.15 shows the measured speed of every valid step, as stored by PROST for each step.

**Figure C.11:**
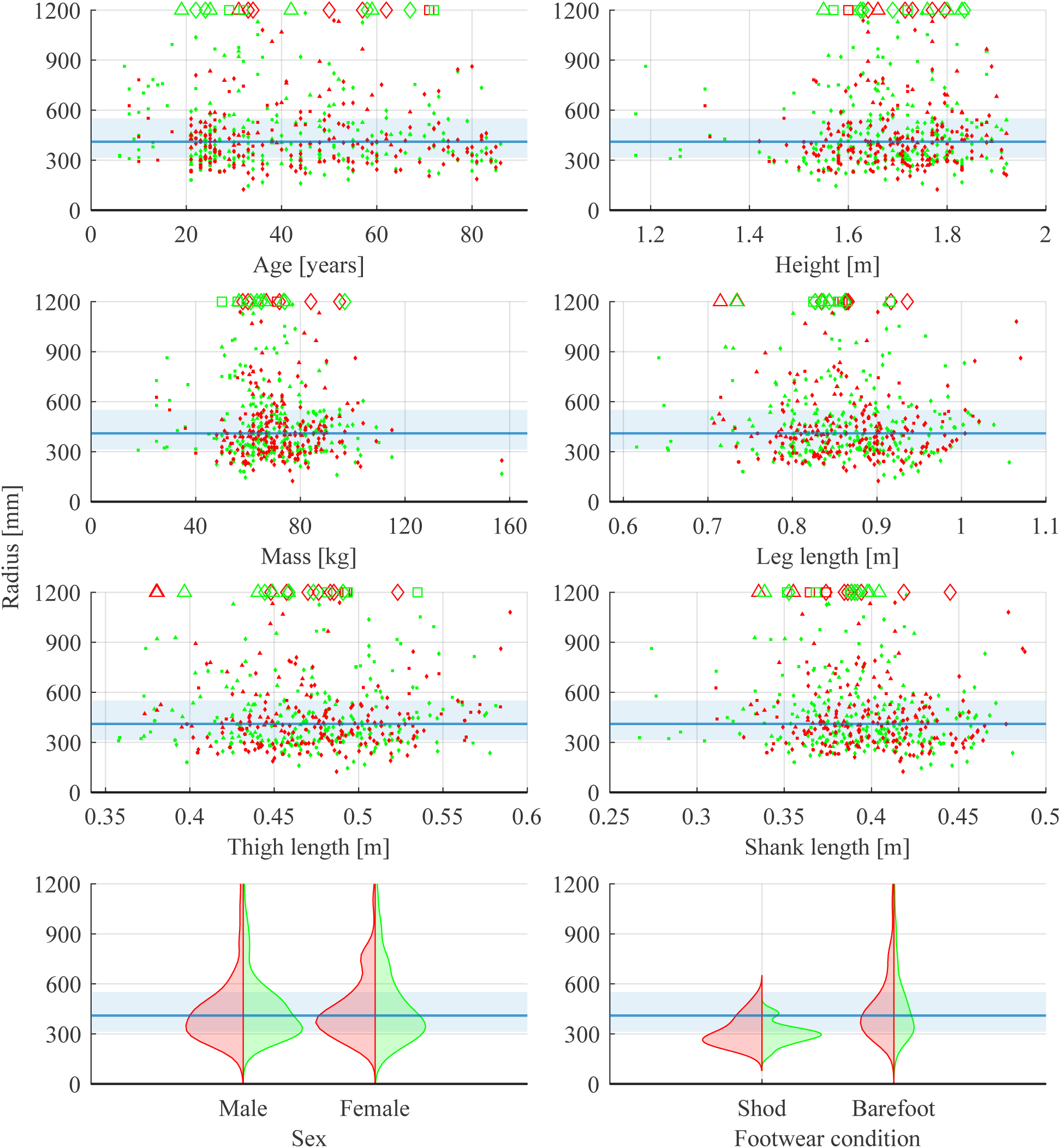
Radius vs all anthropometric parameters and footwear condition for all leg circular arc fits (*N_L_* = 467 legs). Marker shape identifies the dataset (circle, triangle, square, and diamond, for datasets 1, 2, 3, and 4, respectively). Colour identifies the trailing leg (green: right; red: left). For sex and footwear condition, split violin plots show the right-leg (green, right half) and left-leg (red, left half) distributions. The solid blue line and shaded band mark the pooled median and interquartile range; the solid black line marks zero. The y-axis shows at least 95% of legs (below 1200 mm); the 22 legs outside this range are shown as open markers on the upper axis edge. The corresponding statistics are given in Table C.11.

**Figure C.12:**
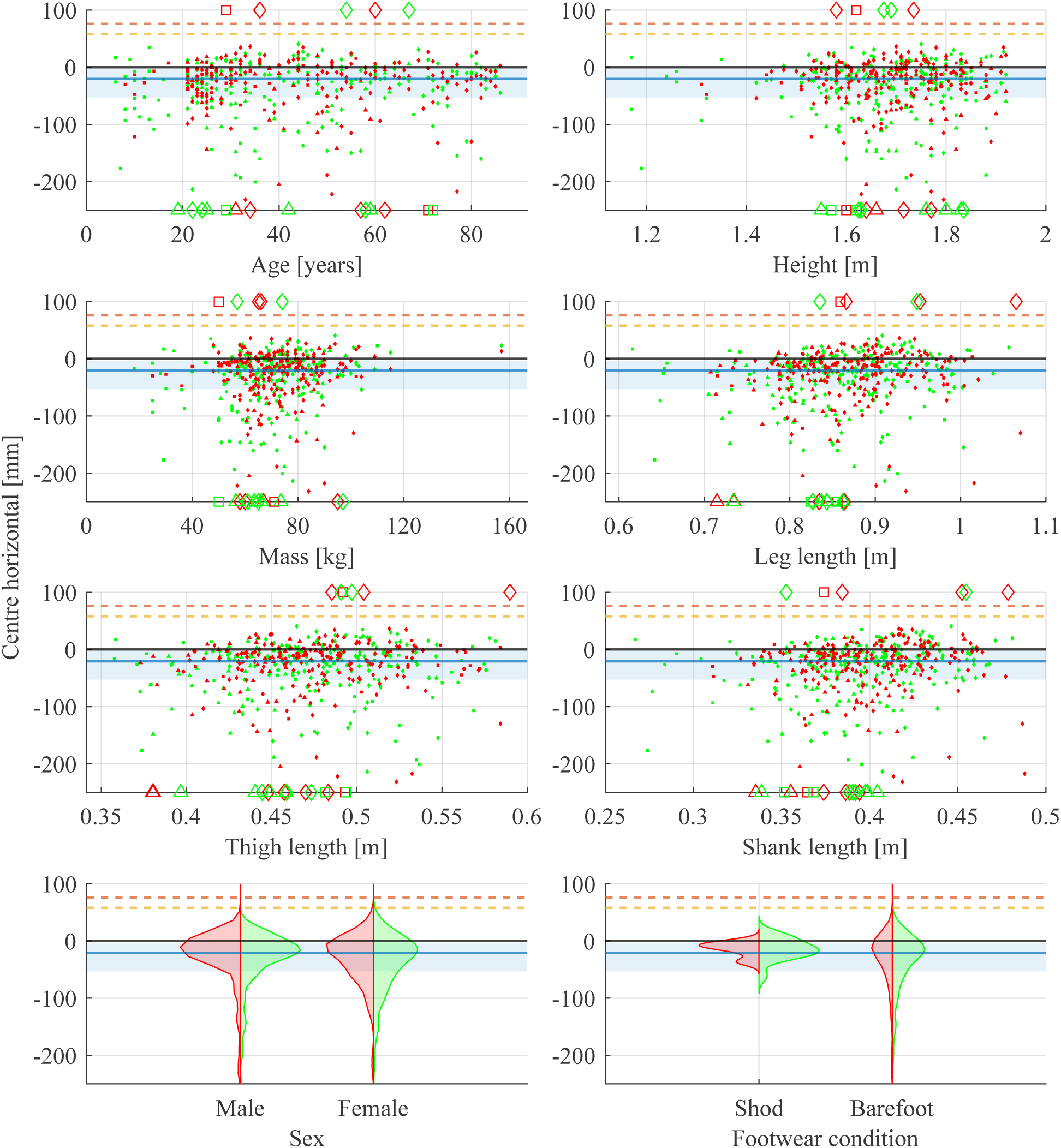
Centre horizontal coordinate vs all anthropometric parameters and footwear condition for all leg circular arc fits (*N_L_*= 467 legs); positive values are anterior to the ankle. Marker shape identifies the dataset (circle, triangle, square, and diamond, for datasets 1, 2, 3, and 4, respectively). Colour identifies the trailing leg (green: right; red: left). For sex and footwear condition, split violin plots show the right-leg (green, right half) and left-leg (red, left half) distributions. The solid blue line and shaded band mark the pooled median and interquartile range; the solid black line marks zero. Dashed lines show the arc-centre offsets anterior to the tibial axis imposed on rigid arc feet by Adamczyk et al. [17] (76 mm, orange) and Adamczyk and Kuo [18] (58 mm, yellow); these are design values rather than measured ROS. The y-axis shows at least 95% of legs (between −250 and 100 mm); the 22 legs outside this range are shown as open markers on the upper (6) or lower (16) axis edge. The corresponding statistics are given in Table C.11.

**Figure C.13:**
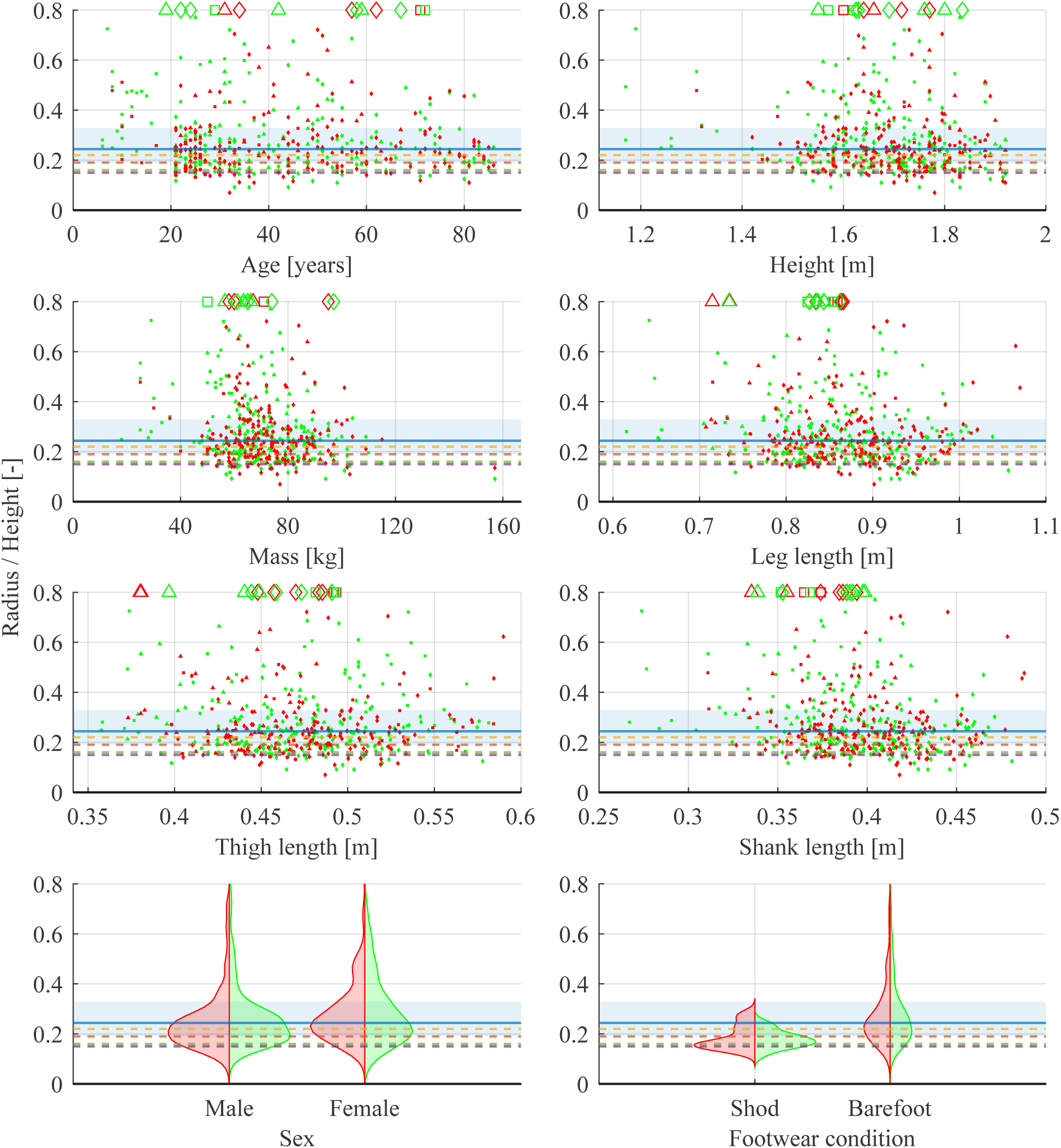
Height-normalised radius vs all anthropometric parameters and footwear condition for all leg circular arc fits (*N_L_* = 467 legs). Marker shape identifies the dataset (circle, triangle, square, and diamond, for datasets 1, 2, 3, and 4, respectively). Colour identifies the trailing leg (green: right; red: left). For sex and footwear condition, split violin plots show the right-leg (green, right half) and left-leg (red, left half) distributions. The solid blue line and shaded band mark the pooled median and interquartile range; the solid black line marks zero. Dashed lines show previously reported height-normalised radii: 0.19 [19] (orange) and 0.22 [20] (yellow) for the ankle-foot ROS, and 0.15 [1] (purple) and 0.16 [21] (grey) for the knee-ankle-foot ROS. The y-axis shows at least 95% of legs (below 0.8); the 18 legs outside this range are shown as open markers on the upper axis edge. The corresponding statistics are given in Table C.11.

**Figure C.14:**
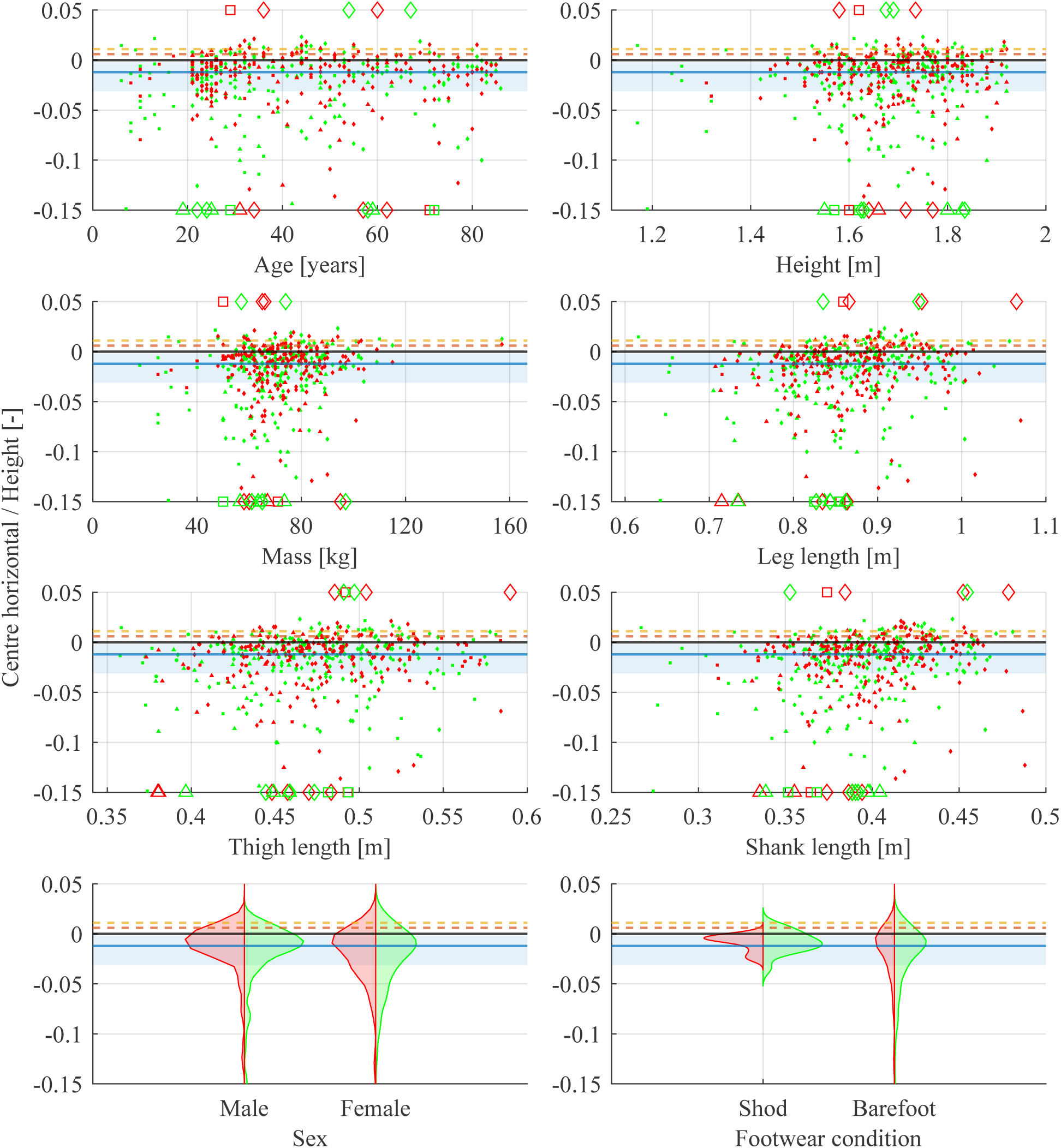
Height-normalised centre horizontal coordinate vs all anthropometric parameters and footwear condition for all leg circular arc fits (*N_L_* = 467 legs); positive values are anterior to the ankle. Marker shape identifies the dataset (circle, triangle, square, and diamond, for datasets 1, 2, 3, and 4, respectively). Colour identifies the trailing leg (green: right; red: left). For sex and footwear condition, split violin plots show the right-leg (green, right half) and left-leg (red, left half) distributions. The solid blue line and shaded band mark the pooled median and interquartile range; the solid black line marks zero. Dashed lines show height-normalised values previously reported for the knee-ankle-foot ROS: 0.006 [1] (orange) and 0.011 [21] (yellow). The y-axis shows at least 95% of legs (between −0.15 and 0.05); the 21 legs outside this range are shown as open markers on the upper (6) or lower (15) axis edge. The corresponding statistics are given in Table C.11.

In Dataset 1, the mean measured speed of each condition was within 0.1 km/h of its nominal value. In Dataset 2, the metronome-cued conditions mostly fell within their nominal bands but overlapped at the band edges, and the self-selected conditions C4 and C5 overlapped C3 and extended up to 8.5 km/h. Dataset 3 covered a wide range (1.1–7.8 km/h) because it included self-paced trials of increasing and decreasing speed in addition to natural-speed trials. Dataset 4, recorded at a single self-selected comfortable speed, had the narrowest distribution. The measured speed ranges of the four datasets therefore overlap only partly, and the datasets differ in speed as well as in protocol.

## Appendix E. Exclusion of Nearly Flat Arcs

A circular arc fitted to a nearly flat CoP path is ill-conditioned: as the arc span *θ* approaches zero, the radius diverges, and a small change in the curvature of the path produces a large change in *r*. For an arc with chord length *L* (the distance between its endpoints),

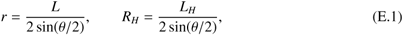

holds exactly, where *L_H_* = *L/H* is the height-normalised chord. The radius is therefore governed by the arc span whenever the chord varies little between legs. Across all *N_L_* = 467 legs, the chord (*L*: median 151.5 mm, IQR 22.8 mm; *L_H_*: median 0.091, IQR 0.016, RCV*_Q_* = 13%) varied far less than *R_H_*(RCV*_Q_* = 43%). Since log(*R_H_*) = log(*L_H_*) − log(2 sin(*θ/*2)), the variance of log(*R_H_*) (0.264) decomposes into that of the chord term (0.021) and that of the arc-span term (0.271), the remainder being their covariance; a single fitted chord (*L_H_* = 0.086) explained 95% of the variance of *R_H_*. Height normalisation does not make the chord more uniform: *L* was only weakly correlated with height (Spearman *ρ* = 0.26), so *L_H_*was weakly negatively correlated with it (*ρ* = −0.25). Median *L_H_* ranged from 0.087 (Dataset 4) to 0.103 (Dataset 1). For the interquartile range of *L_H_*, the cutoff *R_H_* = 1, a fitted radius equal to body height, corresponds to *θ* ≈ 4.8^◦^–5.6^◦^.

The ill-conditioning can also be seen from the sagitta *s* = *r*(1 − cos(*θ/*2)), the largest distance of the arc from its chord. For the excluded legs, the median sagitta was 1.15 mm, about half their *e*_r_, so the departure of the path from a straight line was smaller than the scatter of the points about the fitted arc; across all legs, the median sagitta was 7.1 mm, about four times *e*_r_. This ratio does not by itself separate the excluded legs (63 legs had *s <* 2*e*_r_, 13 of them excluded), and since *e*_r_ is the scatter of the pooled points of all steps, it understates how well the arc of a leg with many steps is resolved.

**Figure D.15:**
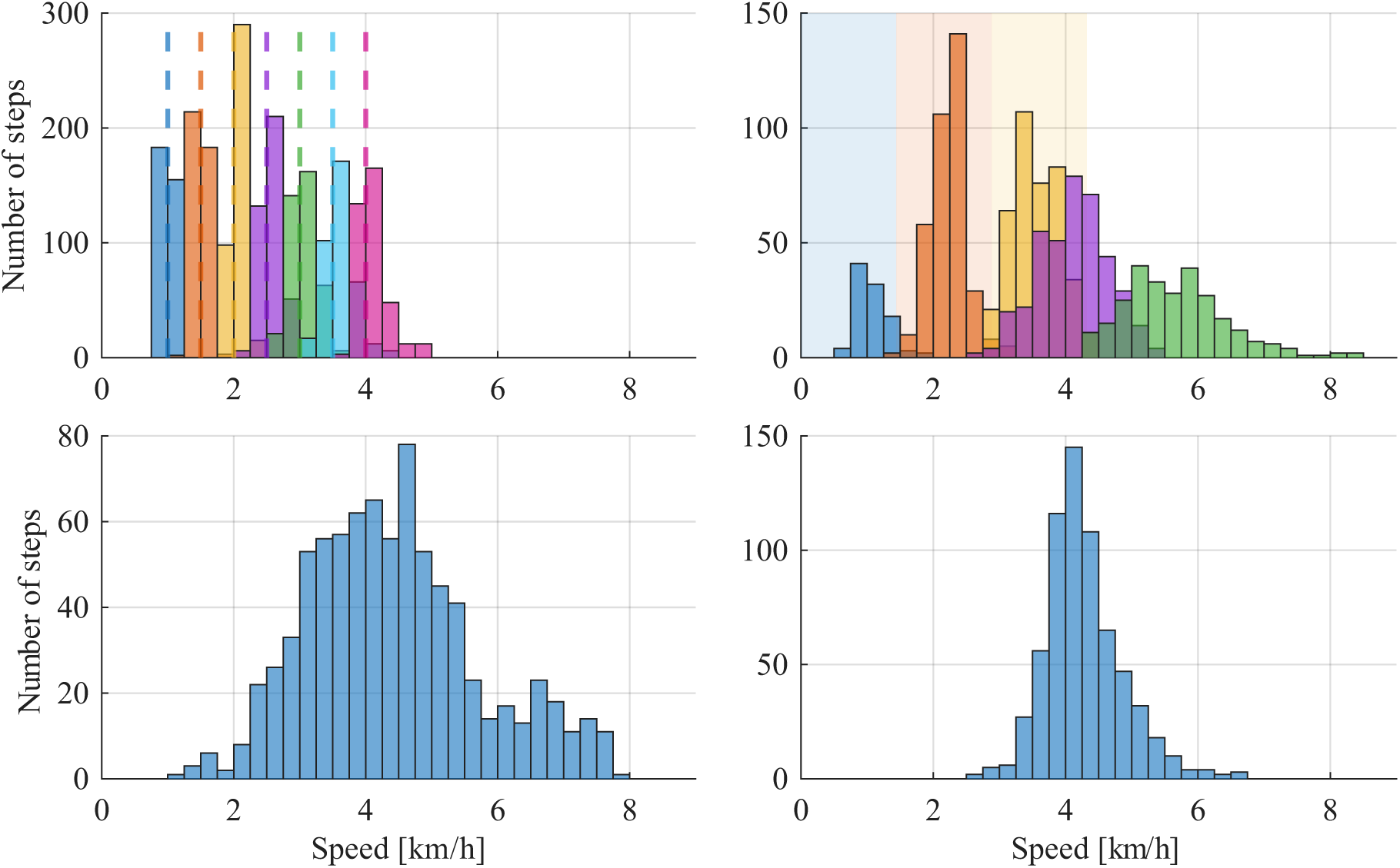
Histograms of the measured walking speed of the valid steps (bin width 0.25 km/h): Dataset 1 [23] (top left), Dataset 2 [24] (top right), Dataset 3 [25] (bottom left) and Dataset 4 [26] (bottom right). Datasets 1 and 2 are shown separately for each speed condition, coloured from the slowest to the fastest condition as blue, orange, yellow, purple, green, cyan and magenta. In Dataset 1, the dashed lines of matching colour mark the nominal metronomecued speeds (1.0–4.0 km/h in steps of 0.5 km/h). In Dataset 2, the shaded regions of matching colour mark the nominal speed bands of the metronome-cued conditions C1–C3 (0–0.4, 0.4–0.8 and 0.8–1.2 m/s); C4 (purple) is the self-selected and C5 (green) the self-selected fast condition.

The cutoff excluded 14 legs (Dataset 1: 0, Dataset 2: 4, Dataset 3: 3, Dataset 4: 7); three participants lost both legs. The arc-span rule equivalent under the fitted chord (*θ <* 4.9^◦^) selected the same 14 legs and one additional leg (*θ* = 4.89^◦^, *R_H_* = 0.95), which was retained. Flat arcs were not an artefact of fits from few steps: the arc span was unrelated to *n*_steps_ (Spearman *ρ* = −0.01, *p* = 0.77), but it was associated with the deviation of *R_H_* from its dataset median (*ρ* = −0.40, *p <* 0.001).

The cutoff is defined, and converted to an arc span, on the height-normalised scale; its effect is reported below for the modelled responses *r* and log *r*. Without the cutoff, the flat arcs dominated the model of *r*: the largest standardised residual (8.0) belonged to a leg with *θ* = 2.2^◦^, and the participants with both legs flat had predicted random effects of 5.6 to 6.0 SD. Because both legs of these participants were flat, they inflated the participant variance *τ*^2^ rather than the residual variance. Excluding them therefore lowered the ICC, from 0.419 to 0.367 for *r* and from 0.483 to 0.423 for log *r*, so the cutoff works against, not in favour of, the reported participant-level variability. On the log scale, the influence of the flat arcs was smaller (largest standardised residual 5.2; random effects up to 4.2 SD).

Table E.12 compares the applied cutoff with arc-span thresholds from 3^◦^ to 20^◦^. The log *r* ICC remained between 0.40 and 0.51 for all thresholds, and the participant random intercept was supported at every threshold tested (boundary-corrected LRT at 5^◦^, 8^◦^ and *R_H_* ≤ 1: *p* ≤ 2×10^−4^ for *r* and *p <* 10^−7^ for log *r*). Normality of the Pearson residuals of the log *r* model was not rejected at the 12^◦^ and 15^◦^ thresholds, which excluded 65 (14%) and 108 legs, and was rejected again at 20^◦^; for the *r* model it was rejected at every threshold except 20^◦^ (200 legs excluded). Since no threshold gave clearly better model behaviour, *R_H_* ≤ 1 was retained as the smallest exclusion that removes the ill-conditioned fits, rather than a threshold chosen to optimise the ICC.

**Table E.12:** Sensitivity of the mixed-effects model to the exclusion of nearly flat arcs. ICCs are from REML fits of the model in Equation 6 to *r* and to log *r*. *A*^2^ (*p*): Anderson–Darling statistic of the Pearson residuals of each model, with its p-value. Under the fitted chord, the applied cutoff *R_H_* ≤ 1 corresponds to *θ* ≥ 4.9^◦^.

| Rule | Legs excluded | *r* |  | log *r* |  |
|---|---|---|---|---|---|
|  |  | ICC | *A*<sup>2</sup> (*p*) | ICC | *A*<sup>2</sup> (*p*) |
| None | 0 | 0.419 | 46 (< 0.001) | 0.483 | 8.6 (< 0.001) |
| *θ* ≥ 3° | 4 | 0.458 | 33 (< 0.001) | 0.481 | 6.7 (< 0.001) |
| *θ* ≥ 5° | 15 | 0.289 | 20 (< 0.001) | 0.396 | 4.2 (< 0.001) |
| *θ* ≥ 8° | 28 | 0.358 | 13 (< 0.001) | 0.442 | 2.5 (< 0.001) |
| *θ* ≥ 10° | 47 | 0.424 | 5.9 (< 0.001) | 0.464 | 0.93 (0.018) |
| *θ* ≥ 12° | 65 | 0.517 | 2.8 (< 0.001) | 0.509 | 0.38 (0.40) |
| *θ* ≥ 15° | 108 | 0.465 | 0.82 (0.034) | 0.440 | 0.72 (0.059) |
| *θ* ≥ 20° | 200 | 0.504 | 0.69 (0.070) | 0.440 | 1.2 (0.003) |
| *R<sub>H</sub>* ≤ 1 | 14 | 0.367 | 19 (< 0.001) | 0.423 | 4.1 (< 0.001) |

**Table F.13:** Coefficients of the covariate models. Each covariate set was added separately to the base model (Equation 6). Estimates are given with 95% case-bootstrap CIs (5000 resamples) as [lower upper]. Side and sex are differences between the two categories. For log *r*, height and mass enter as log *H* and log *M*, so their coefficients are elasticities; *u_c_*coefficients are in mm per unit of the covariate. *v*^∗^: dimensionless walking speed.

| Covariate | Subset | log *r* | *u<sub>c</sub>* (mm) |
|---|---|---|---|
| Side (right−left) | All datasets | 0.008 [-0.046 0.063] | -8.3 [-16.1 -0.7] |
| Side (right−left) | Dataset 4 | -0.044 [-0.11 0.026] | -0.04 [-9.6 9.7] |
| Sex (male−female) | All datasets | -0.092 [-0.20 0.016] | 9.9 [-6.8 28.3] |
| Age (per year) | All datasets | 0.0003 [-0.0023 0.0029] | -0.05 [-0.51 0.39] |
| Height (log *H*; per m) | All datasets | 0.78 [-0.18 1.76] | -20 [-94 53] |
| Mass (log *M*; per kg) | All datasets | -0.087 [-0.38 0.23] | 0.18 [-0.43 0.63] |
| *v*<sup>\*</sup> | Datasets 3–4 | 2.14 [1.14 3.09] | -260 [-455 -78] |
| *v*<sup>\*</sup> | Dataset 4 | 2.55 [1.18 3.78] | -314 [-582 -56] |

## Appendix F. Covariate Models: Coefficients and Sensitivity Analyses

Table F.13 gives the coefficients of the covariate models summarised in Table 8 and Table 10, together with the leg-side effect. Table F.14 and Table F.15 give the variance components of all covariate models, including three checks: leg side, which varies within participants and therefore tests whether the two legs can be pooled rather than explaining between-participant variance; log leg length (or leg length for *u_c_*) in place of height; and Dataset 4 alone, the largest dataset, whose pseudo-*R*^2^ does not average over datasets with different variance components.

**Table F.14:** Variance components of all covariate models of log *r*. Each covariate set is added to the base model of its block, fitted on the same legs, and pseudo-*R*^2^ is computed against it; it is not given for side, which varies within participants and cannot account for between-participant variance. *N_p_/N_L_*: participants/legs. Estimates are given with 95% case-bootstrap CIs (5000 resamples) as [lower upper]. Anthropometrics: age, sex, log *H* and log *M*; with leg length, log leg length replaces log *H*. The Dataset 4 base model is the per-dataset model of Figure 8; its CI differs from the one reported there by up to 0.01 because the two bootstraps draw different resamples.

| Model | *N*<sub>*p*</sub>/*N*<sub>*L*</sub> | *τ*<sup>2</sup> | ICC<sub>*log*</sub> | Pseudo-*R*<sup>2</sup> |
|---|---|---|---|---|
| All datasets: base | 246/453 | 0.066 | 0.42 [0.25 0.57] | – |
| + side | 246/453 | 0.066 | 0.42 [0.25 0.57] | – |
| + anthropometrics | 246/453 | 0.066 | 0.42 [0.24 0.57] | -0.001 [-0.026 0.12] |
| + anthropometrics, leg length | 246/453 | 0.066 | 0.42 [0.24 0.57] | -0.002 [-0.024 0.11] |
| Dataset 4: base | 135/258 | 0.093 | 0.55 [0.35 0.70] | – |
| + side | 135/258 | 0.093 | 0.55 [0.36 0.71] | – |
| + anthropometrics | 135/258 | 0.095 | 0.56 [0.35 0.70] | -0.025 [-0.043 0.12] |
| + *v*<sup>\*</sup> | 135/258 | 0.074 | 0.49 [0.29 0.65] | 0.20 [0.03 0.44] |
| Datasets 3–4: base | 184/331 | 0.085 | 0.52 [0.34 0.67] | – |
| + *v*<sup>\*</sup> | 184/331 | 0.071 | 0.48 [0.29 0.64] | 0.17 [0.02 0.37] |

**Table F.15:** Variance components of all covariate models of *u_c_*. Each covariate set is added to the base model of its block, fitted on the same legs, and pseudo-*R*^2^ is computed against it; it is not given for side, which varies within participants and cannot account for between-participant variance. *N_p_/N_L_*: participants/legs. *τ*^2^ in mm^2^. Estimates are given with 95% case-bootstrap CIs (5000 resamples) as [lower upper]. Anthropometrics: age, sex, height and mass; with leg length, leg length replaces height. The Dataset 4 base model is the per-dataset model of Figure 8; its CI differs from the one reported there by up to 0.01 because the two bootstraps draw different resamples.

| Model | *N<sub>p</sub>*/*N<sub>L</sub>* | *τ*<sup>2</sup> | ICC | Pseudo-*R*<sup>2</sup> |
|---|---|---|---|---|
| All datasets: base | 246/453 | 1649 | 0.47 [0.24 0.65] | – |
| + side | 246/453 | 1647 | 0.47 [0.24 0.66] | – |
| + anthropometrics | 246/453 | 1673 | 0.47 [0.24 0.65] | -0.014 [-0.037 0.078] |
| + anthropometrics, leg length | 246/453 | 1659 | 0.47 [0.23 0.65] | -0.006 [-0.034 0.10] |
| Dataset 4: base | 135/258 | 2133 | 0.57 [0.29 0.77] | – |
| + side | 135/258 | 2125 | 0.57 [0.29 0.77] | – |
| + anthropometrics | 135/258 | 2190 | 0.58 [0.29 0.77] | -0.027 [-0.054 0.097] |
| + *v*<sup>\*</sup> | 135/258 | 1797 | 0.53 [0.26 0.73] | 0.16 [-0.01 0.41] |
| Datasets 3–4: base | 184/331 | 2152 | 0.56 [0.31 0.75] | – |
| + *v*<sup>\*</sup> | 184/331 | 1898 | 0.53 [0.28 0.72] | 0.12 [-0.01 0.32] |

